# Real-time visualization reveals Mycobacterium tuberculosis ESAT-6 disrupts phagosome via fibril-mediated vesiculation

**DOI:** 10.1101/2024.04.19.590309

**Authors:** Debraj Koiri, Mintu Nandi, P M Abik Hameem, Aher Jayesh Bhausaheb, Geetanjali Meher, Assirbad Behura, Akhil Kumar, Vineet Choudhary, Sandeep Choubey, Mohammed Saleem

## Abstract

*Mycobacterium tuberculosis (Mtb)* evades host defense by hijacking and rupturing the phagosome, enabling it to escape to the host cytosol for its survival. ESAT-6, a secreted virulence protein of *Mtb*, is known to be critical for phagosome rupture. However, the mechanism of ESAT-6-mediated disruption of the phagosomal membrane remains unknown. Using *in vitro* reconstitution and numerical simulations, we discover that ESAT-6 polymerization remodels and vesiculates phagosomal membrane. In contrast to the pore formation triggered by a bilayer-spanning conformation, we find that the binding of ESAT-6 to the phagosomal membrane is shallow. Such shallow insertion leads to membrane shape transition leading to tubular and bud-like deformations on the membrane in a concentration-dependent manner, facilitated by the reduction in membrane tension and compressibility modulus. Strikingly, our observations suggest that ESAT-6 polymerizes in bulk and on the membrane, both *in vitro* and in macrophage. Numerical simulations demonstrate that growing fibrils generate both radial and tangential forces causing local remodeling and shape transition of the membrane. Using micropipette aspiration, we quantitatively show that ESAT-6 bound tensed membrane undergoes local changes in membrane curvature and lipid phase separation, also facilitated by the direct contact of the bacteria inside the phagosome. Nonetheless, the vesiculation of the buds is primarily driven by the forces exerted by the polymerization of ESAT-6. Such ESAT-6 mediated vesiculation induces apoptosis and host cell death in a concentration and time-dependent manner that promotes infection. Overall, the findings provide mechanistic insights into the long-standing question of phagosome disruption by *Mtb* for its escape.

## Introduction

*Mycobacterium tuberculosis* (*Mtb*), the causative agent of tuberculosis, survives inside the host by regulating the phagosome maturation and rupture to escape into the cytosol of the host cell [1]. ESAT-6 (6kDa early secretory antigen target), a highly immunogenic effector secreted by *Mtb*, has been identified as a critical protein involved in the process of phagosome (also referred to as Mycobacteria closed vacuole) rupture [2]. ESAT-6 is known to be secreted as ESAT-6: CFP-10 heterodimeric complex, stabilized by two-salt bridges. This is crucial for the secretion and the stability of ESAT-6 in the intracellular environment of the phagosomal lumen [3]. Native ESAT-6/CFP-10 heterodimer extracted from *Mycobacterium tuberculosis* was found to dissociate under acidic pH, however, only ESAT-6 but not CFP-10 is known to interact with the phagosome membrane [4]. Purified recombinant ESAT-6 has been implicated in lysis of artificial lipid membranes [4], red blood cells [5] and cultured macrophages through permeabilization of both the phagosome membrane as well as cell membrane upon gaining access to the cytoplasm [6–8]. Purified ESAT-6, but not other ESX-1-secreted proteins, has been shown to induce pore formation in host cell membranes [5, 9, 10]. The membrane-interacting activity of ESAT-6 is known to largely depend on acidic pH-dependent conformational changes in vitro [9]. It has also been reported that many of the pore-forming activities ascribed to ESAT-6 are due to the residual detergent contamination in ESAT-6 purification adopting widely used protocols [11, 12]. ESAT-6 is found to induce significant deformation of nonacidified phagosome. Yet under acidic conditions liposome membrane is known to undergo lysis induced by both the native as well as recombinant ESAT-6 in the absence of contaminating detergents [13].

Interestingly, a recent study reported that ESAT-6 alone is not sufficient for phagosome disruption. Instead the direct bacterial physical contact was found to be important for gross phagosomal membrane disruption [11]. *In vivo* observations also suggest that ESAT-6 required additional factors such as the lipid phthiocerol dimycocerosate for the phagosomal disruption [12, 14, 15]. *Mycobacterium marinum* transposon mutants have also been found to be capable of phagosome damage despite their inability to secrete ESAT-6 hinting that ESAT-6 alone may not be sufficient for the ESX-1 virulence function [16]. However, there is also strong evidence supporting ESAT-6’s direct role in phagosome lysis and virulence [9, 17, 18]. Most recently, the C terminus of ESAT-6 was shown to be required for phagosomal membrane damage [13]. Though the ability to damage the phagosomal membrane has emerged as central to *Mtb* virulence, the mechanism of ESAT-6 mediated phagosome membrane disruption remains poorly understood [1]. In particular, the insights on the dynamics of ESAT-6 interaction and membrane remodeling of phagosome are lacking[2].

In this work, we demonstrate that ESAT-6 undergoes fibrillation both *in vitro* and *in* vivo, and subsequently disrupts the phagosome through vesiculation. Our findings reveal that the binding of recombinant ESAT-6 to the membrane is shallow, penetrating only to C5 of the acyl chain in the interacting leaflet of the bilayer, rather than spanning the entire bilayer as previously believed. We discover that the ESAT-6 leads to the formation of small tubules to spherical buds on the membrane surface in a concentration-dependent manner, accompanied by a reduction in membrane tension and compressibility modulus. However, bud fission occurs only around 24 hours, coinciding with the timescale in which ESAT-6 surprisingly undergoes fibrillation, forming a network of randomly aligned polymers both in vitro and in vivo. Numerical simulations demonstrate that growing fibrils can generate both radial and tangential forces, causing local remodeling and reshaping of the membrane. We quantitatively show that the binding of the lower oligomer of ESAT-6 on a membrane under tension, as a result of forces generated by fibrils, triggers local changes in membrane curvature and lipid phase separation. The lipid phase separation in the phagosome membrane may also be induced by the direct contact associated with the bacterial load. Finally, the vesiculation of the bud takes place by a three-fold increase in the areal strain on the phagosomal membrane that subsequently induces apoptosis and host cell death likely helping in the infection.

## Result

### ESAT-6 penetrates till C5 of the acyl chain and induces tubular deformations in the phagosome membrane mimic

*M. tuberculosis* blocks phagosomal maturation by secreting the virulence effector protein ESAT-6 through the ESX-1 secretion system. ESAT-6 is known to have a membranolytic activity that helps *M. tuberculosis* escape into the cytosol [1, 13]. However, the mechanism of ESAT-6-mediated disruption of the phagosomal membrane remains unknown. The binding affinity of ESAT-6 on phagosomal membrane mimic (40% DOPC, 5.5% DOPE, 22% SM, and 32.5% Chol)[19] was quantified using the tryptophan intrinsic fluorescence of ESAT-6. Upon titration of 5 μM of ESAT-6 with the increasing lipid concentration (ranging from 0 to 600 μM) using liposomes, we observed a concentration-dependent reduction in the fluorescence intensity of ESAT-6 tryptophan residues hinting at weak association with the membrane (i.e, membrane binding affinity, K_d_ ∼ 400μM; (Fig. 1b). This is in line with the recently reported rapid self-association of ESAT-6 as a result of its low dissociation constant (i.e, K_d_ ∼ 1.5μM) [20]. The decrease in fluorescence intensity signifies that the tryptophan residues are interacting or embedded within the hydrophobic environment of the membrane. The maximum tryptophan fluorescence of ESAT-6 was observed between 315 nm to 335 nm (Extended Data Fig. 1).

**Figure 1.**
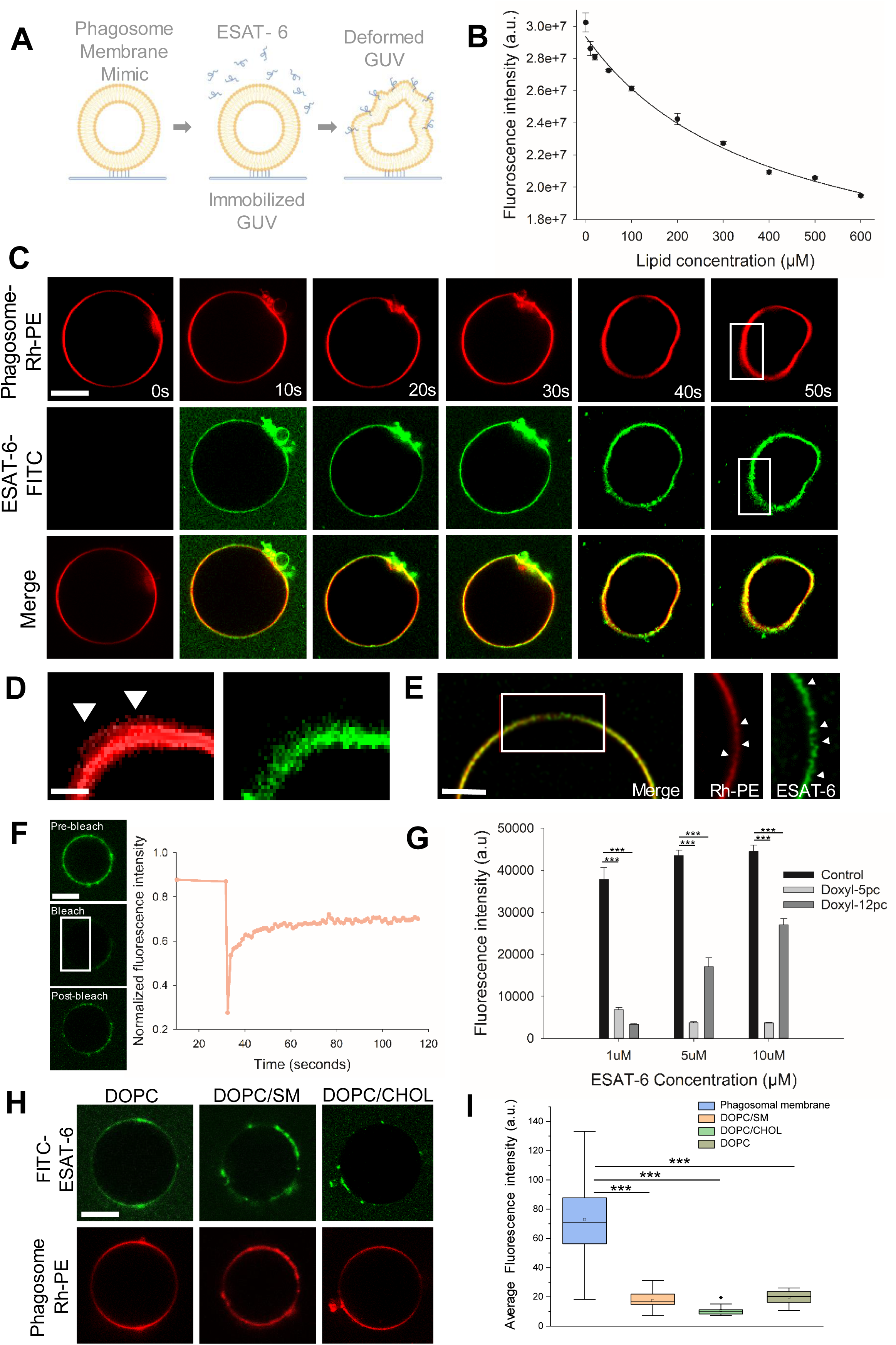
Shallow insertion of ESAT-6 induces changes in phagosomal membrane curvature. **a**, Schematic representation of experimental design depicting deformation of the immobilized GUV mimicking the phagosomal membrane by ESAT-6**. b,** Fluorescence intensity plot of tryptophan residue of ESAT-6 as a function of lipid concentration (LUVs), fitted using hyperbolic decay from sigma plot. Experiments were carried out in PBS pH 5.5 at 25 °C. ESAT-6 concentration was kept constant at 5μM. Data points shown are means ± S.D. of three independent measurements**. c,** Time-lapse confocal imaging of GUVs labeled with 0.1% Rh-PE (red) and immobilized using 0.03% PEG-Biotin incubated with 5μM FITC-ESAT-6 (green). Scale bar, 5μm. Images are the representative of five independent experiments (n=73 GUVs). **d,** zoomed images extracted from **c** represent the curvature change on the membrane. **e,** Structured illumination microscopy (SIM) images of GUVs with the same condition as in **c** followed by zoomed images of the change in membrane curvature. Scale bar, 5μm (n= 15 GUVs). **f,** Mean fluorescence recovery curve after photobleaching experiments of 5μM FITC-ESAT-6 on the membrane. Data points using mean values from three independent experiments. **g,** Fluorescence intensity plot of tryptophan residue of ESAT-6 interaction with LUVs (100nm sizes) containing 10mol% of 5-doxyl-PC and 12-doxyl-PC. Data points shown are means ± S.D. of nine measurements from three independent experiments. **h,** confocal imaging of GUVs of DOPC, DOPC/cholesterol, and DOPC/SM labeled with 0.1% Rh-PE (red) and immobilized using 0.03% PEG-Biotin incubated with 5μM FITC-ESAT-6 (green). Scale bar, 5μm. Images are the representative of five independent experiments (n=25 GUVs). **i,** Mean fluorescence intensity plot of ESAT-6 binding for each membrane condition (***p<0.001 in one-way ANOVA).

To visualize the dynamics of ESAT-6 interaction in real-time, we first immobilized giant unilamellar vesicles (GUVs, 10-50μm) mimicking phagosomal composition doped with 0.01% biotinylated-PEG to facilitate conjugation to streptavidin passivated surface. The GUVs were labeled with 0.1% Rhodamine-PE and incubated with 5μM of FITC-labeled recombinant mycobacterial ESAT-6 (*see methods*). Though this is topologically opposite, the micrometer-scale of the phagosome and the cell membrane will both have a zero mean curvature and thus appear planar for ESAT-6 of 10-20 Å width [21]. ESAT-6 homogenously bound to the GUV surface within 10 seconds of protein injection, followed by tubular deformation of the immobilized GUVs within a minute (Fig. 1c and S1-Movie & Extended Data Fig. 2). Though fine tubular structures were observed on the GUV membrane surface upon the injection of ESAT-6, it is difficult to rule out the possibility that the observed structures arise due to the broad point spread function in the confocal imaging (Fig. 1c-d). Visualizing protein-induced small tubular structures, particularly during their emergence, can be challenging as they are mostly below the optical resolution [22, 23]. The super-resolution image of the equatorial plane of the ESAT-6 bound GUV revealed discrete local binding and changes in curvature capturing the onset of membrane shape transition (Fig. 1e). The diffusion of the membrane-bound ESAT-6 was found to only partially decrease as evident from significant fluorescence recovery after photobleaching, and the rate of diffusion was estimated to be 0.11 μm^2^/sec using the Soumpasis equation as described in the method (Fig. 1f). This also supports the observed weak-association of ESAT-6 with the membrane in the monitored time scale. To examine the ESAT-6 insertion in the membrane, we performed the steady-state fluorescence using depth-sensing fluorescence quenching probes 5-Doxyl-PC & 12-Doxyl-PC with a constant concentration of 10 mol%. Depth-sensitive fluorescence quenching allows estimating the depth of membrane penetration of a fluorophore attached to the target molecule with the quenchers attached at various depths of the membrane bilayer [24, 25]. ESAT-6 has three tryptophan residues in its sequence that allowed us to monitor the fluorescence intensity of the tryptophan in the presence of the doxyl probes (Fig. 1g). In contrast to the bilayer spanning model for pore formation [4, 5, 10, 26], we observed that as the concentrations of ESAT-6 increased, the 5-Doxyl-PC that corresponds to a depth ∼12 Å from the center of the bilayer, showed more quenching effect compared to the 12-Doxyl-PC. This suggests that ESAT-6 does not penetrate the entire bilayer instead it inserts only till C5 of the acyl chain in the interacting leaflet. We further looked at the lipid specificity of ESAT-6 interaction in the phagosomal lipid environment and found that ESAT-6 has a preferential binding for the phosphatidylcholine (DOPC) head group. Amongst the reconstituted minimal lipid membranes of the phagosome model membrane, the highest binding intensity was observed for the DOPC membrane, followed by the DOPC/Sphingomyelin membrane. The presence of 30% cholesterol in the DOPC membrane results in the weakest binding of ESAT-6 hinting at the possible role of packing densities in the binding of ESAT-6 (Fig. 1h). Though the observed difference in binding intensities of ESAT-6 between individual lipid membranes is quite low, however, the phagosomal membrane showed many folds higher binding (Fig. 1i).

### ESAT-6 binding reduces the membrane tension and makes it more compressible

To understand the observed shape elasticity and deformations in the phagosome model membrane induced by ESAT-6, we first examined the fate of the cell membrane tension upon ESAT-6 binding. As the initial composition of the phagosomal membrane is derived from the cell membrane[27], particularly, the outer leaflet of the cell the membrane being the same as the inner leaflet of the engulfing phagosome. We thus monitored the fluorescence lifetime change in the cell membrane of cultured THP-1 differentiated macrophages treated with ESAT-6 over ten minutes and doped with 100nM of Flipper-TR, a membrane tension sensor, [28, 29]. We found that binding of 5μM ESAT-6 to macrophage cell membrane results in a decrease in the fluorescence lifetime from 4.2 ns to 3.4 ns indicating a significant reduction in the cell membrane tension (Fig. 2a, c). It is reasonable to say that ESAT-6 interaction with the phagosomal membrane would also reduce the membrane tension. Indeed, the fluorescence lifetime of the tension sensor in the reconstituted GUVs mimicking phagosomal membrane decreased from 4.5 ns to 3.7 ns upon the interaction with 5μM ESAT-6 (Fig. 2b, c) suggesting a reduction in the tension. The magnitude of the observed changes is significant as previously reported for endosomes [29].

**Figure 2.**
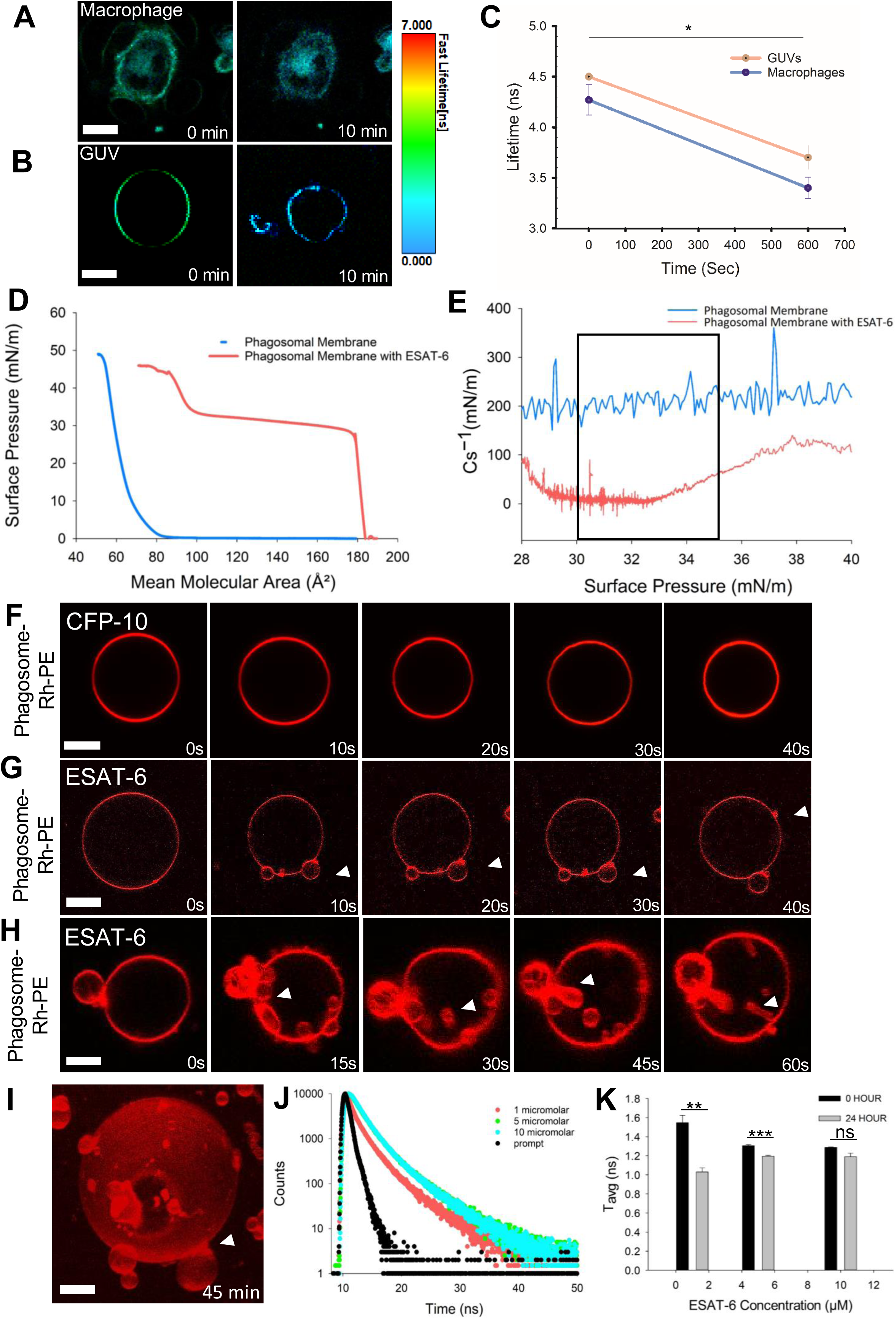
ESAT-6 binding reduces phagosomal membrane tension and compressibility modulus and induces budding at higher surface density. Images from fluorescence lifetime imaging microscopy (FLIM) of FliptR-labelled **a,** live macrophages, and **b,** phagosomal membrane mimic GUV before and after 10 min of 5μM ESAT-6 treatment. **c,** FliptR lifetime measurements of live macrophages and GUV from three independent experiments as shown in **a,b**. the data points shown are means ± S.D (*p<0.05 in one-way ANOVA).**d,** surface pressure (π)– mean molecular area (A) isotherm of the phagosomal membrane in the absence(blue) and presence of ESAT-6 (red). **e,** compressibility moduli (Cs^−1^) at a surface pressure of 30-35mN/m (at the bilayer equivalence pressure marked in the rectangular box) is shown in the graph with the same labeling order and color scheme for the membrane conditions as stated in **d.** All the monolayer experiments were performed on PBS subphase pH 5.5 at 25°C. Each isotherm corresponds to the mean of three independent experiments. Time-lapse confocal imaging of immobilized GUVs (same composition and labeling as in **fig.1c**) in the presence of higher concentration(20μM) **f,** CFP-10 and **g,h,** ESAT-6. Scale bar, 5μm (n=14 GUVs for CFP-10 and n=18 GUVs for ESAT-6). **i,** 3D projection of budded vesicles on the membrane extracted from the **g. j,** Time-resolved fluorescence intensity decay plots of 1 (red), 5 (green), and 10 μM (blue) of ESAT-6, respectively, in the presence of phagosomal membrane mimic LUVs and measured after 24 hours. The fluorescence decay plot of the instrument response function (IRF) is shown in black. All the measurements were carried out in PBS (pH 5.5) at 25°C. These are the representative curves from three independent experiments. **k,** the mean fluorescence lifetime plot extracted from **j** and time-resolved fluorescence intensity decay plots of 1 (red), 5 (green), and 10 μM (blue) of ESAT-6, respectively, in the presence of phagosomal membrane mimic LUVs and measured after 30 minutes (reference, 0 hr), and the data points shown are means ± S.D. of three independent measurements (**p<0.01, ***p<0.001 in one-way ANOVA).

We next examined the response to compression, in the plane of the membrane, upon the binding of ESAT-6 using lipid monolayer membranes. Langmuir monolayers enable quantification of lateral lipid interactions and compressibility modulus [30–32]. First, the surface activity of ESAT-6 was established through its interfacial behavior at the air/buffer interface in the absence of lipids. The addition of 5 μM ESAT-6 in the buffered sub-phase equilibrated to 0 mN/m led to an increase in the surface pressure to 27 mN/m and stabilized over ∼1 hour suggesting the surface active and amphiphilic nature of ESAT-6 (Extended Data Fig. 3). In the presence of phagosomal model lipid monolayer equilibrated at bilayer equivalence pressure of ∼31 mN/m (i.e, bilayer equivalence range of 30-35 mN/m), the addition of ESAT-6 led to the rise in surface pressure up to ∼33 mN/m over an hour hinting at weak interaction with membrane (Extended Data Fig. 4). We next examined the ESAT-6-induced changes in area per lipid molecule and phase behavior of the phagosomal model monolayer membrane. Analysis of the ESAT-6 bound phagosomal model membrane surface pressure (*π*)-area isotherm (*A*) revealed a plateau at 28-32 mN/m suggesting the transition from a liquid-expanded (*l_e_*) to a liquid-condensed (*l_c_*) phase (Fig. 2d). The compressibility modulus (C*_s_*^-1^) of the ESAT-6 bound membrane was extracted from the *π-A* isotherm using the following equation –

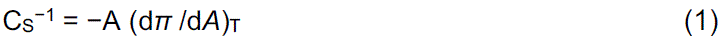

The C*_s_*^-1^ of the monolayer at the bilayer equivalent surface pressure of ∼ 30-35mN/m is considered to be analogous to the elasticity of a bilayer membrane because the hydrophobic free energy and area per lipid are similar to that in membrane bilayer at this pressure [30, 32, 33]. The Cs^-1^ of the phagosomal membrane decreased two-fold from ∼180 mN/m to ∼ 90 mN/m upon the interaction of ESAT-6, suggesting the membrane is more compressible and thus, easier to deform (Fig. 2e).

### Higher surface density of ESAT-6 induces phagosomal membrane budding and initiation of vesiculation

*Mycobacterium tuberculosis (Mtb)* relies on the ESX-1 secretion system for the secretion of ESAT-6 inside the phagosome. The physiological concentration of ESAT-6 inside the phagosome remains unknown, however, considering the confined volume of the phagosome, the local concentration of the secreted protein can be high. Thus, we used a range of concentration from 1-20 μM in line with the previous *in vivo* and *in vitro* studies that used concentrations ranging from 1-50 μM[34–36]. Likely, the effective concentration of the protein inside the confined volume of the phagosome will rise over time. Considering the size of phagosome to be in the order of 2-5 μm, the ratio of the available surface area and number of secreted protein molecules inside a confined volume should be very low. We questioned whether the rise in the phagosome membrane-bound surface density of ESAT-6 induces any further remodeling and shape deformation. To capture this in real-time, we incubated 20 μM of unlabeled ESAT-6 with the immobilized phagosomal membrane mimic GUVs labeled with 0.1% Rhodamine-PE. Extrapolating the change in the area per lipid molecule of a monolayer membrane upon the binding of unlabelled ESAT-6 for a GUV surface, the surface density of the ESAT-6 was estimated to be ∼ 74 x 10^4^ molecules bound on GUV with a radius of 15 μm. (Extended Data Text). To establish the functional role of ESAT-6 binding, we opted not to use FITC-labelled ESAT-6 at higher concentrations to rule out any potential hindrance caused by the FITC molecule on protein binding. Interestingly, time-lapse imaging revealed that ESAT-6 induces large membrane tubules and buds on the membrane having both positive and negative curvature (Fig. 2g-h and S4-Movie). No such deformations were observed when the membrane was treated with other proteins (i.e., 20 μM CFP-10, BSA) at high concentrations ruling out the role of osmolarity changes as a result of protein flux and proving the specificity of ESAT-6 interaction (Fig. 2f and S2-3 Movie). The equatorial plane of the ESAT-6 bound GUVs showed some seemingly detached free vesicles, however, we think most are still connected with tubular necks that remained out of focus. Occasional vesiculation mediated by ESAT-6 crowding cannot be ruled out as observed in the z-stack and such observation was not common. The buds remained attached even after 45 mins of monitoring without undergoing any visible change (Fig. 2i). Size distribution analysis revealed that most of the budded vesicle sizes are less than 3 μm compared to the mother vesicle sizes around 15-20 μm, in diameter (Extended Data Fig. 6). Though the observed size appears to be several folds higher than the intracellular scale, however, we think that the size of the bud might scale with the area available on the mother vesicle and lead to a broad range of bud size as the mother vesicle loses area [37]. We speculate that for a phagosome of size ∼2 μm this might result in the formation of phagosome buds much smaller in size often making it challenging to visualize as well as demarcate from other vesicular compartments due to technical limitations.

Previous reports have shown that Mycobacterial translocation from phagolysosome or phagosome into the cytoplasm, in a non-apoptotic cell, can take upto few hours to days and involves ESAT-6 secretion as well as direct contact of the bacteria[6, 11]. We hypothesized that prior to the *Mtb* escape, ESAT-6-mediated remodeling should reflect in the polarity of the phagosomal membrane interface. Fluorescence lifetime is known to be sensitive to the changes in the local environmental polarity of the membrane [38]. We quantified the fluorescence lifetime of the tryptophan residue of ESAT-6, at 30 min and 24 hours post-treatment, by incubating 1 μM, 5 μM, and 10 μM ESAT-6 concentrations with the phagosomal model membrane. The fluorescence lifetime of the tryptophan residue of ESAT-6 decreased around 24 hours, but not at 30 min, suggesting an increase in the local polarity over time (Fig. 2j & Extended Data Fig. 5). This decrease in the fluorescence lifetime could be due to the insertion of water molecules into the membrane. However, at higher concentrations of ESAT-6, the change in the fluorescence lifetime is less pronounced due to a relatively greater number of proteins on the membrane which hinders the insertion of water molecules (Fig. 2k). Overall, the observations suggest that though tubulation and budding is a fast process, the local polarity increased only at later time scales of ∼ 24 hours till which the predominant number of buds remain undetached.

### ESAT-6 fibrillation drives the rate of vesiculation of the phagosomal membrane

Many of the bacterial secretory proteins have been speculated to possess amyloidogenic propensity[39]. Perplexed by the observation that the buds predominantly remain attached to the phagosomal membrane we next examined ESAT-6 behavior in the presence of a phagosomal membrane interface, for a longer duration of time. Thus, we first investigated whether ESAT-6 shows any fibrillation characteristics estimated through Thioflavin T (ThT) fluorescence assay (Fig. 3a). Surprisingly, we discovered that 20 μM ESAT-6 undergoes rapid fibrillation under the phagosomal acidic pH of ∼ 5.5 without any visible nucleation or lag phase and reaches the equilibrium phase around ∼24 hours, unlike other fibrillating proteins that take much longer[40, 41] (Fig. 3a). Aggregated ESAT-6 structures were also visualized in solution via microscopy around ∼24 hours (Extended Data Fig. 8). On the contrary, the presence of the phagosomal membrane interface leads to a significant slowdown of the ESAT-6 fibrillation kinetics as evident from the nucleation phase lasting around ∼20 hours. Subsequently, fibrillation began around ∼20 hours and reached the equilibrium by ∼35 hours (Fig. 3a). Certain lipids have been found to inhibit or retard the fibrillation kinetics of several amyloidogenic proteins [42] that might explain the observed slowdown in fibrillation kinetics. Further, no fibrillation of CFP-10 was observed in solution or in the presence of membrane confirming the specificity of ESAT-6 fibrillation (Extended Data Fig. 7). The fibrillation behavior of ESAT-6 led us to test whether the same can happen inside THP1-derived macrophages under cellular environment. Indeed, upon incubating recombinant FITC labeled ESAT-6 with THP-1 derived macrophage treated with a NIT-OH (a red fluorescent dye specific for cell membrane), the ESAT-6 is phagocytosed (Fig. 3b). Further, the phagocytosed ESAT-6 vesicles pinch off from the cell membrane and the ESAT-6 intensity inside the phagosome is found increase by 8-fold compared to the extracellular environment (Fig. 3c). Such puncta are often related to the aggregated forms of the proteins as reported earlier for ESAT-6 as well as other amyloidogenic proteins[42–44]. Subsequently, the phagosome numbers decrease over a few hours, suggesting the disruption and cytosolic access of the ESAT-6 (Fig.3d). A striking fibrillar network spanning the entire equatorial plane of the cell was visualized after 24 hours via live-cell imaging through confocal as well as structured illumination microscopy suggesting continued polymerization of the protein (Fig. 3e).

**Figure 3.**
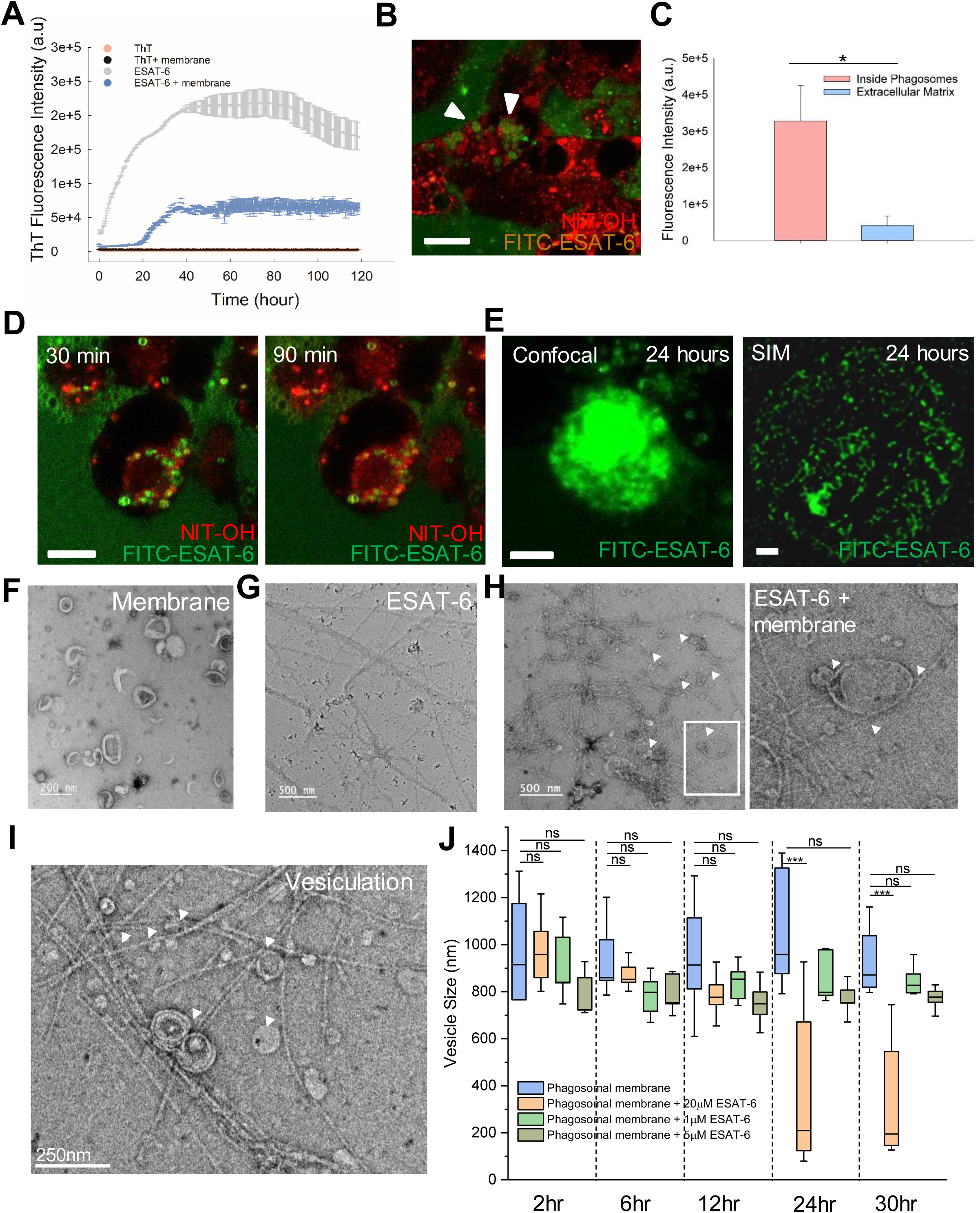
ESAT-6 polymerization facilitates phagosomal membrane vesiculation. **a**, ThT fluorescence assay of only 20μM ESAT-6 (gray) and in the presence of phagosomal membrane mimic LUVs (blue) as a function of time. Data points are shown as the means ± S.D. of three independent measurements. All the experiments are carried out on PBS pH 5.5 at 37°C**. b,** Thp-1 derived macrophages stained with NIT-OH (red) treated with FITC-ESAT-6 (green) showing the biogenesis of ESAT-6 encapsulated phagosomes (highlighted through arrows). Scale bar, 15μm. **c,** Quantification of FITC-ESAT-6 fluorescence intensity through ImageJ of ESAT-6 encapsulated phagosomes (n=150) and extracellular matrix. **d,** Time-lapse imaging of live cell stained by NIT-OH (red) showing ESAT-6 encapsulated phagosomes (green). Scale bar, 15μm. **e,** Confocal images and structured illumination microscope (SIM) images of Thp-1 derived macrophages with 20μM FITC-ESAT-6 incubated for 24 hours. Mesh-like network formed inside cells. Representative images from total number of 87 cells observed in different fields of views. Transmission electron micrographs capturing the shape change of membrane during ESAT-6 fibrillation as shown in **f,** only LUVs (Scale bar, 200nm), **g,** ESAT-6 fibrils (white arrows), and **h,** the budding and vesiculation (white arrows) marked inset images on bottom, **i,** Mesh like network of ESAT-6 fibrils around the mother and daughter vesicles. Scale bar, 500nm. **h,** Vesicle size quantified through dynamic light scattering experiment of only LUVs (blue) and LUVs in the presence of 20 μM (peach),1 μM (green), and 5 μM (dark green) of ESAT-6 monitored for 30 hours. The data points shown are means ± S.D. of three independent measurements (**p<0.01, ***p<0.001 in one-way ANOVA).

We next captured the changes in the phagosomal membrane shape and fibril assembly at ultra-high resolution at the end of the fibrillation phase (i.e., ∼24 hours) using reconstituted large unilamellar vesicles. Negative-stain transmission electron microscopy revealed polymeric fibrillar structures of ESAT-6 both in the presence and absence of the phagosome membrane interface. ESAT-6 fibrils in the absence of membrane seemed to have formed thicker bundles (Fig. 3g). In comparison to the mother vesicle size ranging from 200-300 nm in diameter, most fields of view showed significantly smaller vesicles in the range of ∼ 40-70 nm tangled with ESAT-6 fibrils (Fig. 3h-i and Extended Data Fig. 10). The area of contact of fibrils with the membrane interfaces was found to be both radial and tangential (Fig. 3i). Many of the captured vesicles also showed intact membrane bud necks in contact with the tip of the ESAT-6 fibrils. Quantification of the fibril length revealed most fibrils to be around ∼ 400-800 nm in length (Extended Data Fig. 9). This led us to question whether the ESAT-6 polymerization on the vesicular membrane interfaces is essential for fission. To address this, we used dynamic light scattering to monitor the change of size of mother vesicles in the presence of ESAT-6 over time. Monitoring the reaction containing 20 μM of ESAT-6 and LUVs (ranging ∼ 1 μm in diameter) mimicking the phagosomal membrane, revealed a significant peak shift from 1 μm to ∼70-80nm at 24 hours (Fig. 3j). On the contrary, no significant change in the size was observed for membrane only condition without ESAT-6 (Fig. 3f). Furthermore, both 1 μM & 5 μM of ESAT-6 did not induce any shift in vesicle size suggesting no bud fission over the monitored period of 30 hours (Fig. 3j). The ThT fluorescence kinetics of 1 μM & 5 μM of ESAT-6 on phagosomal model membrane indeed showed a long nucleation phase without any elongation phase confirming the absence of fibrillation (Extended Data Fig. 7). This suggests that ESAT-6 fibrillation is essential for the observed fission of the buds and the kinetics of the fission are dependent on the fibrillation rate that in turn depend on the concentration of the protein.

### The shape transitions of phagosomal membrane deformations are driven by the polymerization forces of ESAT-6

We next examined the role of ESAT-6 fibrillation in inducing membrane deformation and fission. The polymerization of proteins such as actin and β-microglobulin have been reported to generate active forces in driving the transition of a flat membrane to a bud [45, 46]. We used numerical simulations to capture the scaling between the force generated by a growing network of ESAT-6 fibrils and membrane shape. The polymerization of ESAT-6 results in the formation of fibrils with varying lengths (Fig. 3g-h). These fibrils interconnect to create a mesh or a network-like structure typical for polymerizing proteins (Fig. 3h-i). We find that one end of the fibril remains attached to the membrane either radially or tangentially (Fig. 3h-i), and hypothesize that the polymerization at the other end exerts a ratcheting motion against the phagosomal membrane. This ratcheting effect generates forces that play a role in shaping the membrane[34–36]. To model the shape transition of the phagosomal membrane by the ESAT-6 network, we employ Helfrich formalism[47] to account for the curvature-mediated membrane energetics. We explicitly incorporate additional terms in the Helfrich free energy to account for the forces exerted by the fibrils. Helfrich free energy has been widely used to understand membrane surface deformation [46, 48–53].

In our model, the forces exerted by the fibrils include both vertical and radial components (Fig. 4a). The deformed membrane is assumed to exhibit axis-symmetry along the Z-direction (Fig. 3i). Furthermore, we model the double-layered structure as a monolayered surface, serving as a plausible mimic of the phagosomal membrane. The total energy of the membrane (*E*) is defined as

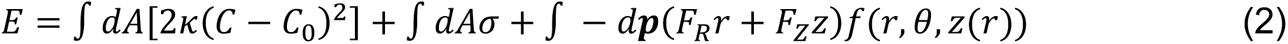

**Figure 4.**
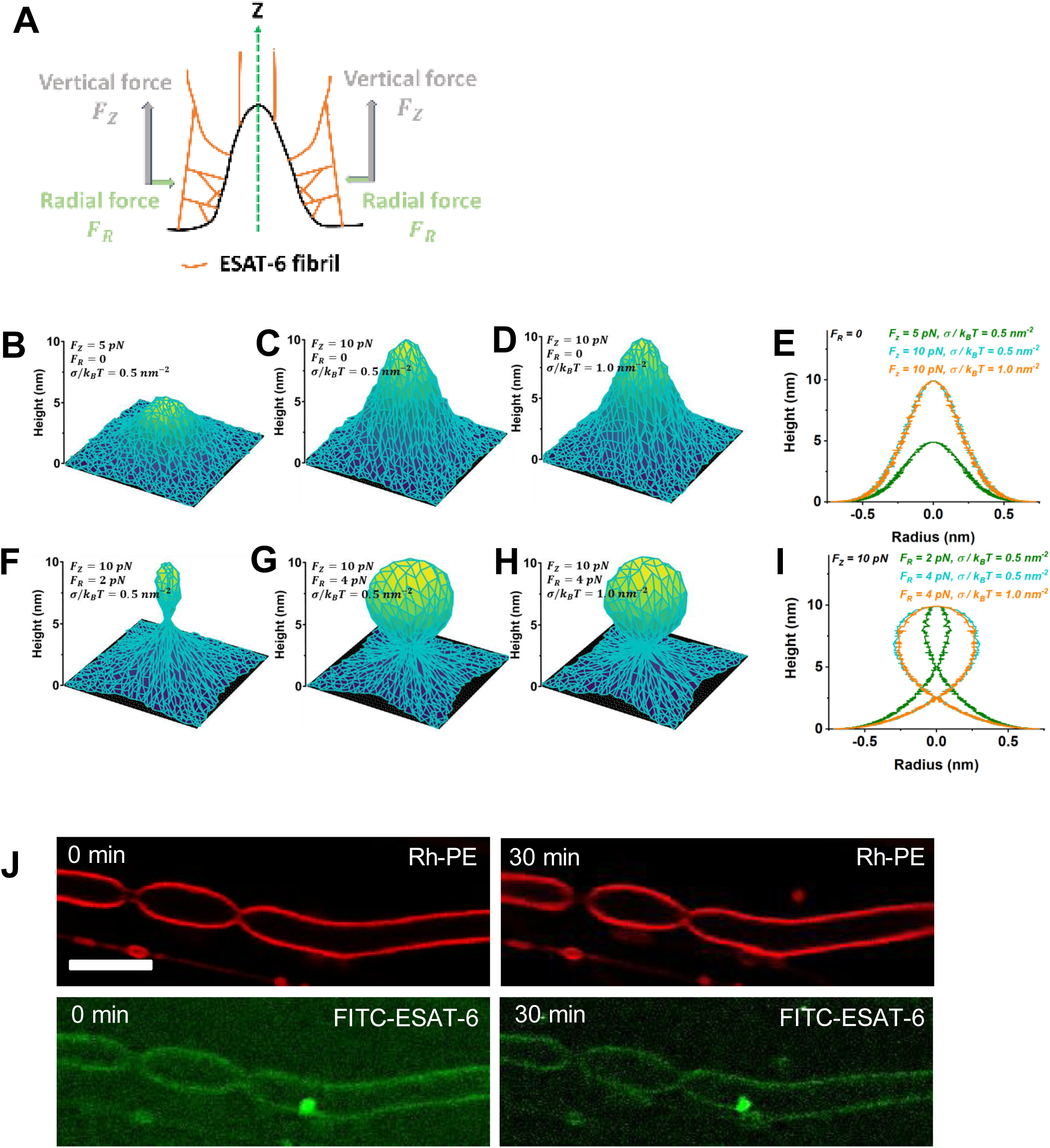
ESAT-6 polymerization forces induce fission. **a**, schematic showing the projections of radial and tangential forces generated by ESAT-6 polymerization. **b,c** A tubular bud emerges with the application of lateral force, F_Z_, while the radial force F*_R_* = The surface tension σ is kept at 0.5*k_B_T*. **d,** when an elevated surface tension (σ = 1.0*k_B_T*) is applied, there are no significant changes in the shape of the tubular bud, even with forces F*_Z_* = 10 *p*N and F*_R_* = 0. **e,** Two-dimensional representation of the bud growth process depicted in **b-c. f,g** The radial force introduction results in a more spherical bud shape, with a connecting neck forming between the bud and the membrane. This effect is highlighted under a constant lateral force F*_Z_* = 10 *p*N. **h,** even with an elevated surface tension (σ = 1.0*k_B_T*), the shape of the spherical bud remains relatively unchanged when subjected to forces F*_Z_* = 10 *p*N and F*_R_* = 4 *p*N. **i,** Two-dimensional representation of radially influenced bud shape shown in **f-h. j,** Time-lapse confocal imaging of free-floating membrane tubes composed of phagosomal membrane composition labeled with 0.1% Rh-PE (red) and 5μM FITC-ESAT-6 (green). Representative image from three independent biological replicates.

The first term in Eq. (2) represents the Helfrich bending energy functional, where *C* stands for the mean curvature of the surface, *C*_0_ for the intrinsic curvature, κ for the bending rigidity, and *dA* for infinitesimal area element of the membrane. The second term defines the energy due to surface tension where σ represents the surface tension of the membrane. The last term accounts for the energy due to forces exerted by ESAT-6 fibrils, where F_R_ and F_*Z*_ represent the radial and vertical forces, respectively. In the last expression, ᵖ = (r, θ, *Z*(r)), and *f*7r, θ, *Z*(r)8 = 1 for the region of the membrane where the fibrillation force is acting; otherwise, *f*7r, θ, *Z*(r)8 = 0 [46]. We minimize the free energy of the system by numerically solving Eq. (2) using Monte-Carlo simulated annealing [46, 53]. For details of the simulations, see Extended Data Text.

Because the binding of ESAT-6 in the phagosomal membrane is shallow, we expect that such binding would induce a localized intrinsic curvature at the binding sites. This leads to an observable deformation in the shape of the membrane (Extended Data Fig. 12). Additionally, we expect that as the surface density of ESAT-6 increases, the number of membrane-bound ESAT-6 fibrils will also rise significantly, resulting in a higher degree of deformation. We chose the vertical and radial forces in the order of pN to mimic biological forces as has been measured for actin and other polymerizing protein networks[34–36]. Systematically applying vertical forces (F_*Z*_) induces changes in the membrane shape, leading to a noticeable increase in the size of the tubular bud (Fig. 4b-e). Furthermore, when radial forces (F_R_) are introduced, a distinct neck forms along with a spherical bud (Fig. 3f-i). This highlights the significance of both vertical (F_*Z*_) and radial (F_R_) forces in reshaping the deformed membrane. Specifically, the vertical force regulates the size of the bud, while the radial force governs the shape, leading to the formation of a constricted neck. Our theoretical analysis suggests that the primary driver of membrane deformation is the forces generated due to ESAT-6 fibrillation. The outcome of this force-induced effect is the formation of a tubular bud, transitioning into a spherical bud, accompanied by the development of a distinct neck. Importantly, this specific shape configuration is crucial for facilitating the subsequent fission process, where the spherical bud detaches from the neck. EM images (Fig. 3h-i) show the co-occurrence of ESAT-6 fibrils meshed with the membrane and the corresponding deformation, thereby supporting our theory of fibrillation-driven membrane deformation. Additionally, the mere presence of free-standing tubular necks without tension and strain did not undergo ESAT-6 mediated fission of the buds (Fig. 4j), further confirming the fibrillation mediated fission.

### ESAT-6 fibrils facilitate membrane fission by exerting forces that increase areal strain and induce local lipid sorting

To validate the observations from the numerical simulations, we subjected the membrane to stretching forces under equiosmolar conditions to rule out any potential osmotic issues. Capturing the force exerted on the membrane by the polymerizing ESAT-6 is technically limiting as fibrillation starts only after nucleation lag period of ∼ 20 hours *in vitro*. To circumvent this, a negative pressure of 50 Pascals that corresponds to 0.5pN/µm^2^ was applied to the phagosomal membrane GUV by micropipette aspiration to mimic the exerted polymerization forces. Biological forces generally operate in the order of pN as has been measured for actin and other fast polymerizing protein networks[34–36]. A stable aspirated tongue length was obtained in the case of the phagosomal membrane GUV that did not undergo any change in the tongue length for the monitored duration of ∼5 minutes (Fig. 5a & S5 – Movie). Upon the injection of the unlabelled monomeric/lower oligomeric ESAT-6, the aspirated tongue length gradually increased followed by an expanding lipid phase separation area in the middle of the tongue (Fig. 5b & S6-Movie). The fluid lipids were found to preferentially partition towards both the termini of the aspirated tongue as evident from the enhanced Rhod-PE fluorescence. Rhod-PE is known to preferentially partition into liquid disordered regions of the membrane resulting in enhanced fluorescence intensity upon lipid segregation[54, 55]. As a result, a high curvature tubular neck formed in the middle of the tongue before the fission and release of the budded vesicle. Upon fission of the tip of the tongue, the remaining tongue relaxed back to the GUV surface and retracted out of the micropipette. Using the Laplace law, we then quantified the change in membrane tension induced by the ESAT-6, through the relationship between surface tension and the pressure difference (Equation 3):

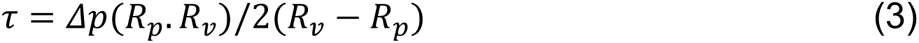

where R_*v*_ is the radius of the vesicle and R_*p*_ is the protrusion radius. Furthermore, estimating the areal strain (i.e, area stretch modulus, Δ*a*/*a*_0_) exerted by the ESAT-6 interaction on a phagosomal membrane under high membrane tension using the equation 4,

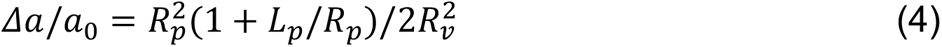

revealed that there is a three-fold increase in the areal strain before the fission of the bud (Fig. 5c & S6-Movie). Analysis of the relationship between change in tension and areal strain revealed that while areal strain increased throughout the fission process, the membrane tension increased up to the constriction point followed by relaxation and attaining equilibrium after the vesiculation. The observed relative change in area of the vesicle as a result ESAT-6 binding rules out any possibility of osmosis[56] (Fig. 5d). We next questioned whether the observed lipid phase separation under high membrane tension is solely a consequence of stretching or could it also be induced by the biochemical exchange of material between the bacteria and phagosomal membrane under physiological conditions[57]. Direct contact of the Mycobacteria with the phagosomal membrane is also considered important for phagosomal rupture in addition to ESAT-6 [11]. To address this in real-time, we first reconstituted *in vitro* live GFP-*Mycobacterium smegmatis* encapsulated inside a phagosomal membrane GUV. The viability of the encapsulated bacteria was found to be 75 % by estimating the colony-forming units by bacteria subjected to electro-formation conditions (Extended Data Fig. 11). The encapsulated bacteria in direct-contact likely induced surface adhesion of the GUV membrane lipids as evident from the merged fluorescence signal at the area of contact (Fig. 5e). The direct contact also resulted in the local bending of the membrane captured in z-stack (Fig. 5f). Additionally, a higher load of the encapsulated bacteria inside the phagosome membrane GUV resulted in both shape deformation as well as significant lipid phase separation wherein the Rhod-PE was observed to preferentially enrich below the equatorial plane of the GUV (Fig. 5g-h, time-lapse & z stack). Reconstituting phagosomal membrane mimic GUV containing the lipids isolated from the mycobacterial cell envelope induced striking phase-separated domains as evident from the preferential partitioning of Rhod-PE (Fig. 5i). This suggests that material exchange as a result of direct-contact cannot be ruled out as one of the potential factors inducing phase separation.

**Figure 5.**
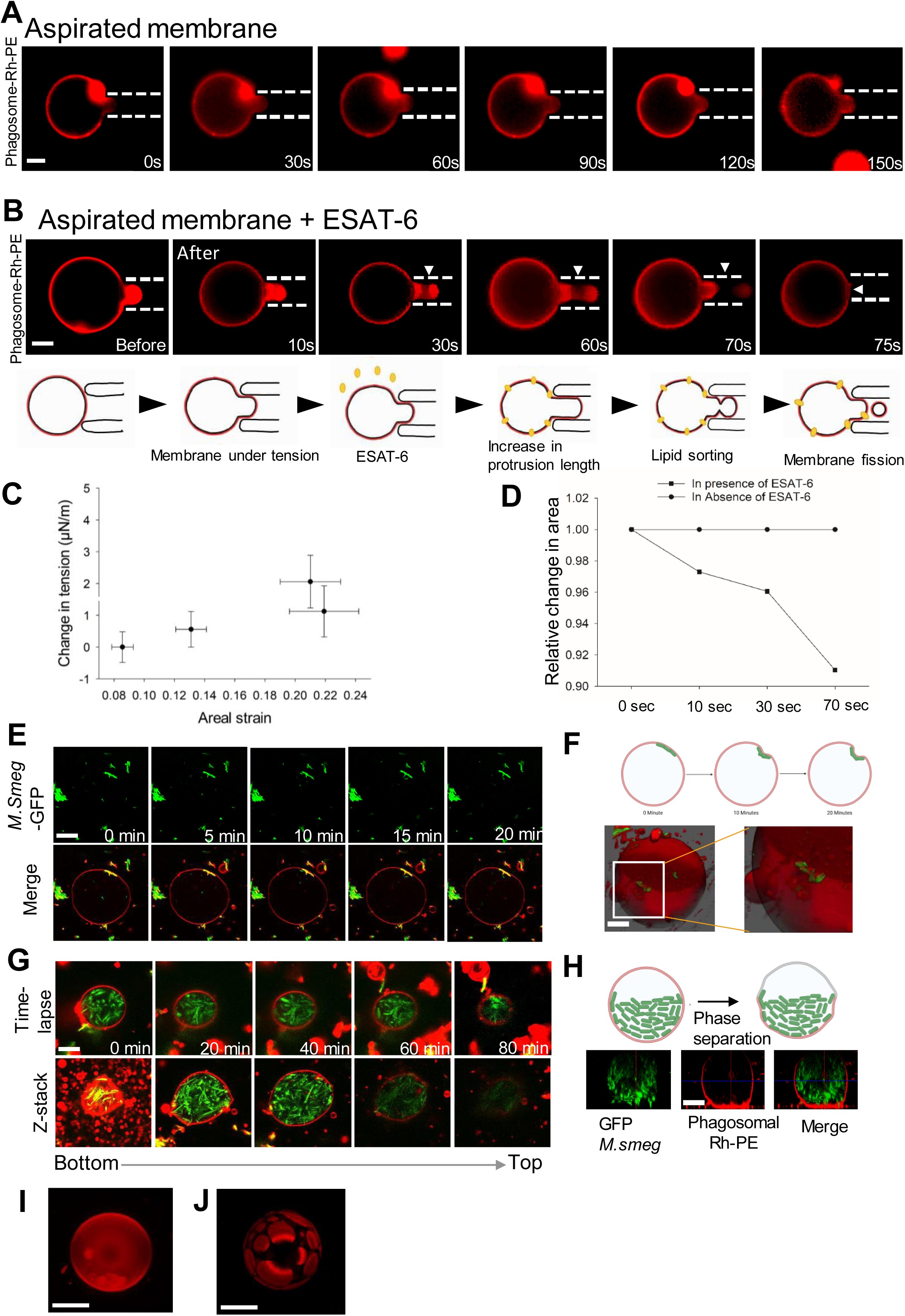
Phagosomal membrane vesiculation proceeds through local modulation of curvature, lipid segregation and an increase in areal strain. **a**, Time-lapse fluorescence imaging of GUVs labeled with 0.1% Rh-PE (red) aspirated with 50pa force and monitored over 150 seconds. **b,** Time-lapse fluorescence imaging of GUVs labeled with 0.1% Rh-PE (red) aspirated with 50pa force followed by the addition of unlabelled ESAT-6 and monitored over 75 seconds Scale bar, 5μm. **a&b** Images are the representative of three independent experiments. **c,** Areal strain vs change in tension plot from **b. d,** Relative change in membrane area plot from **a,b. e,** Time-lapse confocal imaging of GUVs labeled with 0.1% Rh-PE (red) and immobilized using 0.03% PEG-Biotin encapsulated with live *M.smegmatis* (green). Scale bar, 5μm. Images are the representative of three independent experiments. **f,** schematic and 3D projection of the mycobacteria-mediated curvature change from **e. g,** Time-lapse confocal imaging and z-stack of GUVs labeled with 0.1% Rh-PE (red) and immobilized using 0.03% PEG-Biotin encapsulated with the higher load of live *M.smegmatis* (green). Scale bar, 5μm. Images are the representative of three independent experiments. **h,** schematic and z-stack projection of the mycobacteria-induced membrane phase separation of the membrane from **g.** Z-stack projection of **i,** phagosomal membrane (Scale bar 10μm), and **j,** phase separation induced by the lipid mixing of the phagosomal membrane and the mycobacterial cell envelope at 1:1 ratio (Scale bar 5μm) and labeled with 0.1% Rh-PE.

The observed membrane vesiculation *in vitro* as well as the fibrillation of ESAT-6 inside macrophage cells then led us to question the fate of the homeostatic state of the cell. In line with the previous reports [8, 12, 58, 59], we observe that ESAT-6 induced a high rate of apoptosis as evident from the Annexin V-PI staining in a concentration and time-dependent manner, which correlates with the observed in-vitro fibrillation kinetics of the ESAT-6. (Fig. 6a). The high-resolution electron microscopy captured significant morphological changes in the macrophage resulting in the disintegration of the nucleus and cell membrane rupture which leads to the release of cellular contents when treated with 20μM ESAT-6 and incubated for 24 hours (Fig.6b&c). One of the peculiar observations was that most of the cells in different fields of view appeared elongated in shape. This suggests that after the vesiculation of the phagosomal membrane, ESAT-6 escapes into the cytosol and induces apoptosis leading to cell death.

**Figure 6.**
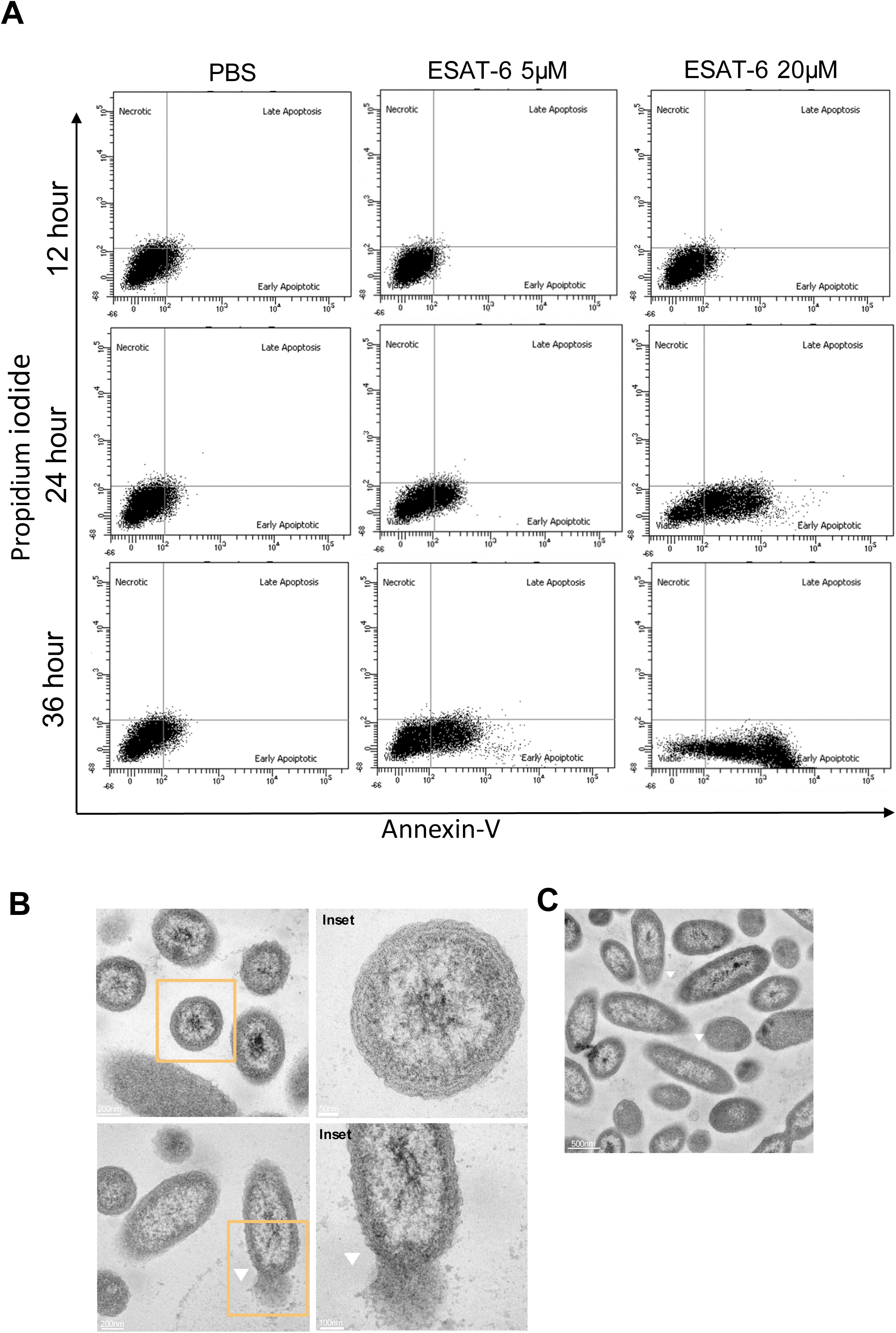
ESAT-6 induced apoptosis and cell death in a concentration and time-dependent manner. **a**, Apoptosis was detected by flow cytometry in macrophage cells after stimulation with ESAT-6 (varying concentration) at different time points. Lower left (double negative), live cells; lower right (annexin V positive, PI-negative), early apoptosis; upper right (double positive), late apoptosis; upper left (annexin V negative, PI-positive), necrosis. Transmission electron micrographs capturing **b,** normal cell, and apoptotic cells are highlighted in the inset. **c,** elongated cells with ruptured membranes shown through white arrows. All the cells were treated with 20μM ESAT-6 followed by incubation for 24 hours. These are the representative images from three independent experiments.

Together, we demonstrate that the initial binding of ESAT-6 induces tubular and bud-like deformations by the reduction in the membrane tension. Both ESAT-6 binding and direct contact with Mycobacterium induced lipid segregation in the membrane which might be important for the rupture of the phagosome. The forces exerted by the ESAT-6 polymerization increase the membrane tension accompanied by the rise in areal strain that drives the vesiculation of the phagosomal membrane, subsequently, inducing apoptosis and host cell death.

## Discussion

In this study, we investigated the mechanism by which the *Mtb* secretory protein, ESAT-6, deforms the phagosome membrane for the escape and survival of the bacteria. We demonstrate that the binding of ESAT-6 on the phagosome membrane mimic is concomitant with the reduction in the overall membrane tension that facilitates the vesiculation of the phagosome membrane in a fibril-dependent manner. While the ESAT-6 is sufficient to induce membrane budding in its lower oligomeric form, the vesiculation of the membrane buds predominantly occurs only upon the polymerization of ESAT-6 into fibrils. The phagosome disruption is one of the key aspects in the arrest of the phagosome maturation and the egress of Mycobacteria [2, 60]. It is reported that *Mtb* ruptures the phagosome membrane and escapes after 48 hours through an ESAT-6-mediated phagosome lysis [6, 7]. ESAT-6, the major secretory protein of ESX-A is membrane-lytic that mediates phagosome rupture, however, the molecular mechanism remains elusive [2, 11, 13, 60]. Though the oligomeric state of the ESAT-6 complex is yet to be determined, it is reported to insert as a membrane-spanning protein that provides the foundation for pore formation based on haemolysis assay [5, 9, 10]. An important reason behind the observed membrane-spanning pore-forming conformation could be the usage of oversimplified lipid composition, that is, DOPC. The packing density arising due to the cylindrical geometry of the DOPC may facilitate the deep insertion of the ESAT-6 [61] but might not be the case for a complex multi-component biological membrane [42]. On the contrary, we show that ESAT-6 inserts only until C5 of the acyl chain, on biologically relevant phagosomal membrane mimic comprising lipid composition of complex geometry. Pore formation may occur transiently as an intermediate step, owing to weak binding to the membrane as confirmed by high K_d_ as well as diffusion (Fig.1b, f). Unlike the ESX-E/F secretory proteins and other pore forming proteins, there is no concrete experimental evidence for pore formation by the oligomeric assembly of ESAT-6 [62–65]. Indeed, the changes in the conductance pattern induced by ESAT-6 measured on artificial lipid bilayers suggest a non-uniform fluctuation and was speculated due to the amyloidogenic properties of ESAT-6 as also seen in some bacterial toxins [26, 39]. ESAT-6 binding to membrane at low and high surface densities induced shape transition from slightly curved to tubular structures (Fig 1c-d) and membrane buds. The binding of the protein makes the phagosome membrane more deformable enabling the generation of curved protrusions as evident from the ESAT-6-induced reduction in the compressibility modulus of the phagosome and membrane tension (Fig. 2e, Fig. 2a-c). However, the buds do not detach from the GUV surface until 24 hours (Fig 2g-h). Further, an increase in polarity was observed over 24 hours resulting from the larger proximity of deformed interfacial regions to water moieties quantified by the lifetime changes of Tryptophan residues of ESAT-6 (Fig. 2j-k). The observed tubulations and buds can be attributed to the changes in local membrane curvature and lateral pressure introduced as a result of high surface density and shallow insertion of ESAT-6 pushing away the lipid-head groups in the interacting membrane leaflet[66–69]. Indeed the magnitude of the protein-induced membrane bending and curvature generation is known to non-monotonously depend on the insertion depth of the protein for biologically relevant conditions [68, 70]. Shallow embedding of amphipathic helices has been shown to induce formation of tubes of varying sizes. For example, while epsin and synaptotagmin forms narrow ∼ 20nm diameter tubes [71, 72], N-BAR containing endophilin and amphiphysin generate tubes ranging ∼ 35-50nm in diameter that subsequently form vesicles of the same size [73]. Recently, tubulation and budding was reported in the phagolysosomes containing *E. coli* important for lysosomal reformation during phagosome resolution [74].

We then wondered why the detachment of the membrane buds takes around 24 hours (Fig 2i). ESAT-6 is known to form amyloid-like structures in the inclusion bodies of *E. coli* making the kinetics of transition much faster [39]. However, very recently ESAT-6 has been shown to undergo rapid self-association though lacking any evidence of fibrillation [43]. ESAT-6 indeed gets phagocytosed upon treatment with THP-1 derived macrophage, and subsequently fibrillates and deforms the phagosomal membrane. Further, upon gaining access to the cytosol it continues to polymerize into a strong network of fibrils spreading throughout the cell (Fig 3e). We provide the first evidence supporting fibrillation of ESAT-6 inside the macrophage phagosome. The decrease in the number of phagosome as a result of elevated concentration of ESAT-6 inside the phagosome suggests that the kinetics of the disruption may vary *in vivo* and *in vitro,* depending on the local concentration of the protein. Interestingly, we observe that ESAT-6 fibrillates in the presence of phagosome membrane interface that slows down the fibrillation kinetics and forms a mesh-like network at the equilibrium phase (Fig 3a-c, 3h-i). It is important to note that until the equilibrium phase is reached, ESAT-6 would exist as a mixture of monomers, oligomers, and fibrils, all of which can bind to the membrane with different affinities and deform the membrane [42, 44]. Both the high surface density of the protein and the growing network of polymerizing protein are known to exert forces on the membranes resulting in local modulation of curvature and tension [45, 75–78]. However, lowering the concentration of ESAT-6 to slow down or abort the polymerisation showed no vesiculation, indicating that polymerisation is essential for vesiculation. Keeping the membrane under elevated tension through aspiration to mimic the fibril-mediated force generation, binding of oligomeric ESAT-6 triggers membrane fission accompanied by an increase in the membrane tension and areal strain (Fig. 5b). This might be facilitated by both direct bacterial contact and curvature-driven lipid sorting aided by ESAT-6, as the phagosomal lipid composition is close to the demixing point also observed for endocytic proteins [79, 80]. We propose that the fibrillation of ESAT-6 inside the phagosome might induce vesiculation of the membrane resulting in decrease in the surface area/volume ratio (Figure 7). And this persistent vesiculation eventually induces apoptosis, thus aiding the mycobacterial escape and spread to other neighbouring cells promoting infection. The role of apoptosis during *M.tb* infection is debated [81–85], however, by modulating the concentration of the secreted ESAT-6 inside the phagosome, *M.tb* may regulate its own residency time. The proposed mechanism explains the variation in the phagosomal escape time observed for Mycobacteria and could be crucial for the pathogenesis.

**Figure 7.**
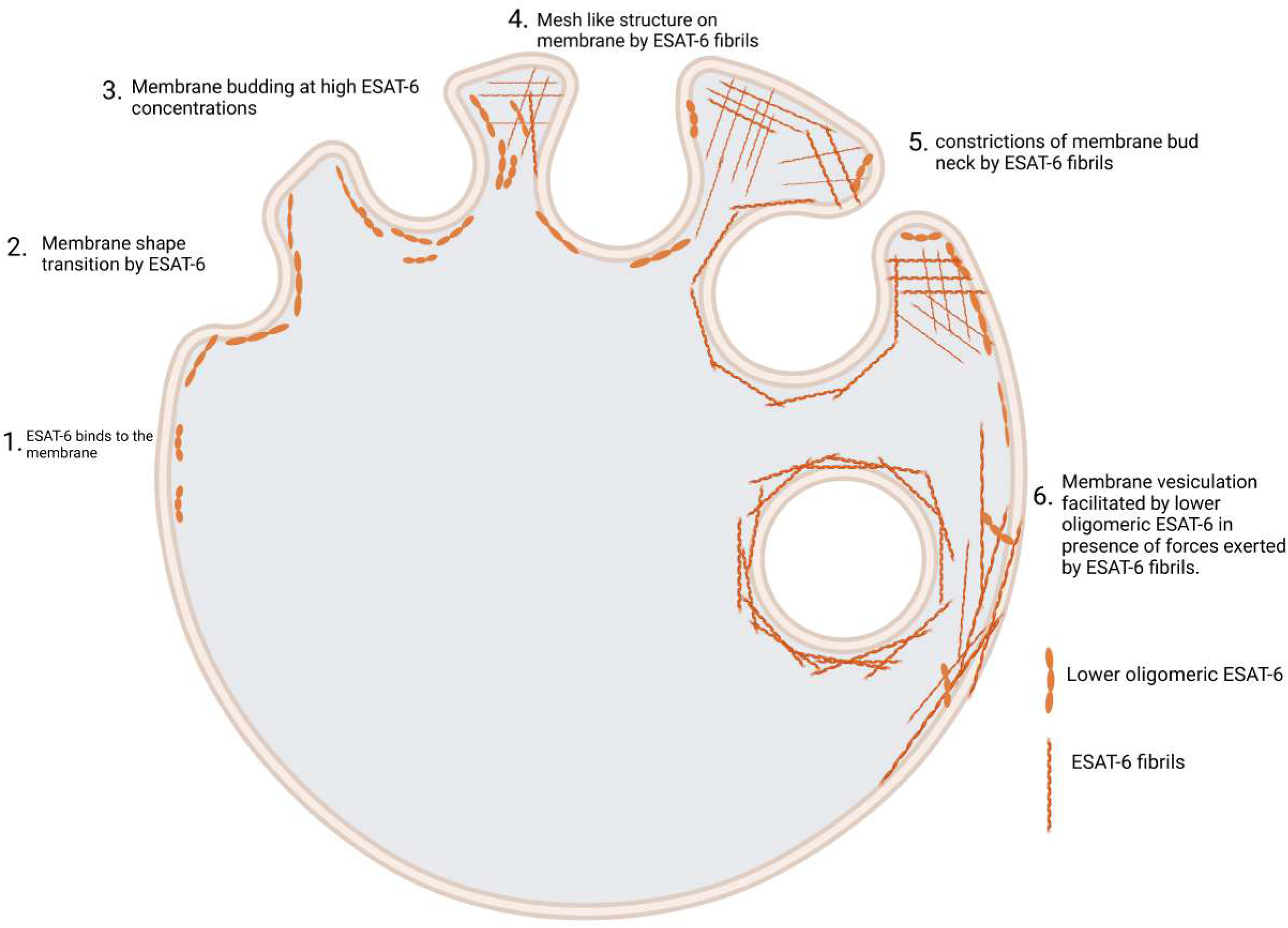
Schematic representation of the proposed mechanism of the ESAT-6 fibril-mediated phagosomal deformation.

## Methods

### Materials

1,2-dioleoyl-*sn*-glycero-3-phosphocholine (DOPC), 1,2-dioleoyl-*sn*-glycero-3-phosphoethanolamine (DOPE), BSM, di-stearoyl phosphatidyl ethanolamine-PEG (2000)-biotin and cholesterol were purchased from Avanti Polar Lipids. Fluorescent conjugated lipid lissamine rhodamine B sulfonyl (Liss Rhod PE) was also purchased from Avanti Polar Lipids. Avidin coated on the chamber slides was purchased from Sigma-Aldrich. ESAT-6 was labeled using the FITC labeling kit purchased from Sigma-Aldrich and labeled using the standard protocol. Proteins were aliquoted in PBS (1X) and stored at –80°C.

### Preparation of GUV

The phagosomal membrane mimic GUVs were composed of 40% DOPC, 5.5% DOPE, 22%BSM, 32.5% cholesterol, 0.03% PEG-Biotin, and 0.1% Rhod PE. To achieve a satisfactory yield of GUVs, the electroformation was performed as described in [23]. In summary, a uniform coat of 15μL of lipid mix at 5 mg ml^−1^ was first dried on conductive indium-tin oxide-coated glass (Nanion technologies, GmbH) under vacuum for at least 1 h. The lipid film was then rehydrated in a PBS (pH 5.5) at 300 ±5 mOsm and GUVs were allowed to grow for 3 h under a sine voltage (2V, 10 Hz) at 60°C. For encapsulation of the live Mycobacteria inside GUVs, the lipid film was rehydrated using 7H9 media with precultured Mycobacteria of OD 1-1.2.

### Overexpression and purification of ESAT-6

E. coli BL21 (DE3, plysS) strain was utilized for the expression and purification of ESAT-6. The strain was transformed with the plasmid pET22b, which included the M. tb gene Rv3875 (protein ESAT-6). The transformed bacteria were grown in LB medium at 37 °C, and protein expression was done at OD600 nm∼0.6, induced by adding 1 mM of isopropyl –D galactopyranoside (IPTG) overnight at 18 °C. Following, bacterial cells were pelleted, and sonicated, and the (His)6-tagged protein was purified using affinity chromatography with Nickel nitrilotriacetic acid (Ni-NTA) according to the manufacturer’s recommendations (GE Healthcare, UK). SDS-PAGE analysis validated the purification of rESAT-6. The purified fractions were dialyzed, concentrated, and kept in aliquots at –80 °C until further use.

### Fluorescence quenching assay

The interaction of ESAT-6 with the phagosomal membrane was observed through the intrinsic fluorescence of Tryptophan residue in ESAT-6. To measure the fluorescence intensity 5μM of ESAT-6 was equilibrated for 5 minutes with increasing lipid concentrations. The excitation wavelength was fixed at 290 nm and the emission spectra were collected between 300-400 nm at a constant temperature of 25 °C. All the fluorescence measurements were performed with an FLS 1000 (Edinburg Instruments, UK) spectrophotometer using a quartz cuvette with a 1cm path length.

### Light Microscopy

8 Well chamber slide from Ibidi was coated with 10μl of 1mg/ml streptavidin, and 200μL of biotinylated GUVs was added and incubated for 15 mins for the GUVs to be immobilized. The final concentration of 5μM of FITC-ESAT-6 was added into the chamber. Imaging was performed on Leica TCS SP8 LCSM using an appropriate laser for Rhodamine-PE (561nm) and Alexa-488 (488nm). Identical laser power and gain settings were used during all experiments.

For the Fluorescence recovery after photobleaching (FRAP) measurements on the GUVs, the GUVs were doped with 0.1% of rhodamine PE and were incubated with 5 μM of ESAT-6. Initially images were captured at reduced laser intensity. Photobleaching was performed at full laser intensity for 30 seconds, resulting in the bleaching within the selected ROI of radius of 5μm. Subsequently, the laser intensity was reverted to the attenuated level and the recovery curve was recorded.The photobleaching was performed at the equatorial plane of the GUV being visualized. The FRAP curves for each condition were repeated three times and then normalized. The diffusion coefficient was calculated using the Soumpasis equation for 2D diffusion

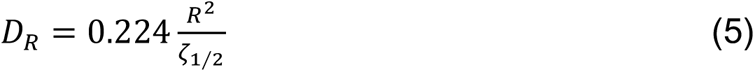

where 0.244 is the numerically determined value, r (5 μm) stands for the radius of the laser beam focused on the region of interest, and τ1/2 is the time required for half the recovery. The time for half the recovery was determined by plotting the normalized recovery curve.

For the in-vivo experiments, the human monocyte cell line (THP-1) was incubated overnight with PMA along with the RPMI to differentiate into the macrophages. Cells were labeled with NIT-OH, a red fluorescent labelling dye. Followed by the treatment with the 20μM FITC-ESAT-6 on the cells and super-resolution images were taken after 24 hours of incubation with the 488nm laser on Zeiss Elyra7.

### Fluorescence Lifetime Imaging

Fluorescence lifetime imaging microscopy was performed on an MT200 (Pico Quant, Berlin, Germany) time-resolved fluorescence confocal microscope with a Time-Correlated Single Photon Counting (TCSPC) unit. The laser power was adjusted between 10 to 100nW for 488nm laser light measured after the dichroic mirror. The Well glass bottom slide was directly placed on immersion water of 60x objective. A dichroic mirror of 488nm was used as the main beam splitter. Out-of-focus emission light was blocked by a 50 μm pinhole and the in-focus emission light was then split by a 50/50 beam splitter into two detection paths. Single Photon Avalanche Diodes (SPADs) served as detectors. Both data acquisition and analysis were performed on the commercially available software Sympho Time 64 (Pico Quant GmbH, Berlin, Germany).

### Time resolved fluorescence measurements

Time-resolved fluorescence intensity decays were obtained by using IBH 5000F Nano LED equipment (Horiba, Edison, NJ) in the time-correlated single photon counting (TCSPC) mode. The excitation source of a pulsed light-emitting diode (LED) was used. At 290nm optical pulses were generated by this LED with a pulse duration of 1.2 ns and emission was measured at 335nm with 1 MHz repetition rate. The Instrument Response Function (IRF) was quantified at the respective excitation wavelength using Ludox (colloidal silica) as a scatterer. 10,000 photon counts were collected in the peak channel to optimize the signal-to-noise ratio. The lifetime of tryptophan residue of ESAT-6 was performed using emission slits with a bandpass of 8 nm. Data were examined using DAS 6.2 software (Horiba, Edison, NJ). Fluorescence intensity decay curves were developed with the IRF and measured using:

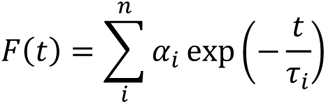

All the plots were quantified with a random deviation of about zero with a maximum χ^2^ value of 1.2 or less. Intensity averaged mean lifetimes τ*_avg_* for fluorescence exponential decays were obtained from the decay times and pre-exponential factors using the following equation

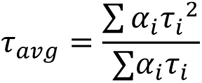

where α_I_ shows the fraction of τ*_i_* lifetime.

### Thioflavin T Assay to determine kinetics of protein aggregation

The aggregation kinetics of ESAT-6 was monitored by ThT fluorescence as a function of time. The final monomeric ESAT-6 concentration was 20μM and 50μM was diluted in PBS solution (Ph-5.5) which comprises 200μM of LUV and 50μM of ThT dye. All the samples were incubated at 37°C in NUNC 96 well plate throughout the experiment. The plate was loaded into the 96-well Clariostar microplate reader equipped with 448/482-nm excitation/emission filters and the ThT fluorescence intensity was monitored every 15 minutes of 500 cycles.

### Lipid Monolayer experiments

The monolayer experiments were performed in KSV NIMA Langmuir balance. The instrument consisted of two barriers used during the compression isotherms assays and as a surface pressure sensor a Wilhelmy microbalance with filter paper plate was used. Monolayers of DOPC: DOPE:SM: CHOL were formed by spreading the lipid/chloroform solution (1mg/ml) in a dropwise manner and kept for 15 mins to equilibrate the monolayers and for the evaporation of the chloroform. PBS (Ph-5.5) was used as the subphase with the maintained temperature of 25^◦^C for all the Langmuir experiments. The final working concentration of 5μM ESAT-6 was injected into the monolayer for isotherm assays. Isotherms were recorded with uniform compression of the monolayer at a constant speed of 1 mm/min. Isotherms were recorded until collapse pressure (πc) was reached. For constant area assays (Time v/s surface Pressure), the monolayer is initially compressed to 30mN/m and allowed to equilibrate for 15 minutes. ESAT-6 was then injected into the subphase and the change in the surface pressure over time was recorded. For all the Langmuir experiments, the trough was cleaned thoroughly with ethanol and filtered double-deionized water until zero surface pressure was reached. Each experiment was performed thrice.

### Negative Staining & Transmission Electron Microscopy

10μl of the fibrillated ESAT-6 with LUVs was incubated in the carbon-coated grid for 20 minutes and wicked off with the help of qualitative filter paper from Himedia. Then 10μl of 2.5% glutaraldehyde was added to the grid for fixation of the sample. The carbon-coated grid was then rinsed using deionized water and 10μl of uranyl acetate was added. After the incubation of 3 minutes and rinsing the grid with deionized water. The grid was then vacuum-dried overnight. The imaging was performed on a JEOL (JEM 2100F) transmission electron microscope. For the cell-based experiments, ThP1-derived macrophages were treated with Esat-6 and incubated for 24 hours. Cells were washed with 0.1M phosphate buffer to get rid of cell culture media and processed for EM as described previously [86]. Briefly, cells were fixed with primary fixative (2.5% glutaraldehyde, 1.25% paraformaldehyde, and 0.1 M phosphate buffer, pH 7.0) for 15 min at RT. Cells were harvested and again resuspended in 1 ml fresh fixative media and incubated on ice for 1 hour. Cells were washed with 0.1M phosphate buffer followed by post-fixation with 1% osmium tetroxide (OsO_4_) in 0.1 M phosphate buffer for 1 hour at RT followed by washing with distilled water twice. Next, cells were incubated with 1% uranyl acetate for 1 hour at RT. Subsequently, samples were dehydrated by incubating in increasing concentrations of ethanol (30%, 50%, 70%, 80%, 90%, 95%, and 100%) for 10 minutes each, with two more incubations in 100% ethanol from a freshly opened bottle. The samples were subsequently embedded stepwise using Spurr’s low-viscosity resin (EMS). Samples were infiltrated for 2 h each with a 3:1, 1:1, 1:3 ethanol / embedding media mixture. Cells were incubated overnight with 100% fresh resin. The next day, cells were again resuspended in fresh 100% resin for 2–3 h, transferred into BEEM capsules (EMS), and polymerized at 70°C for 4 days. Semi– and ultrathin sections were produced with a diamond knife (Diatome) on an ultramicrotome (Ultracut UCT; Leica Microsystems), collected on 200 mesh copper grids (EMS), poststained with uranyl acetate and lead citrate, and visualized with a Talos S200 transmission electron microscope (TEM; Thermo Fisher Scientific), operating at 200 kV. Pictures were recorded on a below-mounted 4k × 4k BM-Ceta (CMOS) camera.

### Micropipette Aspiration

All the measurements were performed on an Olympus IX83 epifluorescence microscope equipped with the Hoffman modulation optics and observed on the 100X objective lens. For positioning the pipette three-axis hydraulic micro-manipulators are used. Micro-pipettes are pulled (Sutter Instruments CO P-87, Narishige Micro forge MF-900) from 10μm diameter capillaries. 100 μl of GUVs were added to the open homemade coverslip. The desired suction pressure for aspirating a GUV is provided, causing the vesicle to be aspirated through the micropipette tip and develop an inner protrusion. Over the period video was recorded to determine the vesicle geometry on the addition of the ESAT-6.

### Subcellular fractionation and lipid extraction of Mycobacterial cell envelope

The *M. smegmatis* were grown in 7H9 broth under continuous shaking for 3 days at 37° C. For subcellular fractionation, the culture was centrifuged at 4400g to obtain the bacterial pellet and carried out following the procedure outlined in [87]Briefly, the bacterial pellet was subjected to mechanical disruption twice using a French press followed by the bead beater by using microbeads (Himedia) to break the cells. Furthermore, the cells were lysed 2-3 times with beads and then centrifuged at low spin to remove unbroken cells. The supernatant was centrifuged 3-5 times at 27000g for 40 minutes to pellet the cell wall. The supernatant obtained previously was then ultracentrifuged at 100,000×g for 1 hour to obtain the cell membrane in the pellet fraction. Finally, the cell wall and cell membrane fractions were resuspended in PBS at a 1:1 ratio and stored in aliquots at –80°C.

Lipid extraction from the mycobacterial cell envelope was done by using the Folch method as described in[88]. The cell envelope was dried with nitrogen gas and kept in a vacuum overnight. The dried cell envelope was mixed with chloroform and then sonicated for 30 minutes at a constant pulse. The solution was again dried and chloroform and ice-cold methanol were subsequently added to the 2:1(V/V) ratio to the sample. After incubation in ice for 20 minutes same amount of distilled water was added as methanol and kept in ice for 10 minutes followed by centrifugation at 1500rpm for 5 minutes to separate the upper polar part containing salt and proteins and the lower organic phase containing lipids. The lower organic phase was dried using nitrogen gas and stored at –80°C.

### Image & Statistical Analysis

The ImageJ software was used to analyze and process images. The fluorescent intensity plots along the equatorial plane of GUVs were quantified using the average fluorescence intensity of the membrane (F_vesicle_) by subtracting the background intensity (F_background_). A uniform detection setting was applied for all sets of experiments. Vesicle sizes were quantified manually using ImageJ. Statistical significance was calculated through one-way ANOVA in sigmaplot where p-values less than 0.05 were considered statistically significant throughout all the experiments.

## Acknowledgements

We thank Thierry Soldati for critical comments on the manuscript. We thank Rohan Dhiman for the kind gift of the ESAT-6 and CFP-10 plasmid and GFP-*M.smegmatis* strain. We thank Subhasis Chattopadhyay for his assistance with the Flow cytometry. We acknowledge all the members of M.S. lab for critical reading of the manuscript. MS greatfully acknowledge the DBT-Wellcome Trust India Alliance (Grant-IA/I/20/2/505212), Department of Atomic Energy for financial support for intramural support and SERB, India for National postdoctoral Fellowship to MN (PDF/2022/001807).

## Notes

### Competing Interest Statement

The authors have declared no competing interest.

### Summary of Updates

1.) Figure 3b is updated showing the phagocytosis of ESAT-6 inside the phagosomes. 2.) Figure 3c Quantification of the fluorescence intensity of the ESAT-6 between phagosome and extracellular matrix. 3.) Figure 3d live cell imaging of ESAT-6 encapsulated phagosomal disruption. 4.) ESAT-6 fibrils spread throughout the cell after 24 hours observed under confocal and super-resolution microscopy. 5.) Figure 4j shows the importance of ESAT-6 polymerization forces in facilitating the fission.

