## Supplementary data text for "Real-time visualization reveals Mycobacterium tuberculosis ESAT-6 disrupts phagosome via fibril-mediated vesiculation"

### Extended data text

### Estimation of the number of ESAT-6 molecules that interacted with the membrane

From figure (2.d), we observed that at 35mN/m surface pressure (π) the area per molecule of the phagosomal membrane is 58 $Å^2$ and the phagosomal membrane in the presence of 5μM of ESAT-6 (molecular weight 11,000 Daltons) is 96 $Å^2$. However, the area of the monolayer trough is 360 $cm^{2,}$ and the volume of buffer in the monolayer trough is 120ml.

Concentration in moles

$$= \frac{Concentration\ of\ protein\ \left(\frac{mg}{ml}\right)}{Molecular\ weight\ of\ protein}$$

$= 5*10^{-6}$ molecules per ml

Number of molecules

$$= \ Concentration\ in\ moles * volume\ of\ monolayer * Avagadro's\ number$$

$= 36*10^{19}$ molecules

Protein density on the trough,

$$= \frac{Number\ of\ molecules}{Area\ of\ monolayer\ trough}$$

$= 10^{18}$ molecules per $cm^2$

Area occupied by protein molecules = (96-58) $Å^2$ = 38 $Å^2$ = $3.8*10^{-15}$ $cm^2$

Number of protein molecules inserted into the monolayer membrane

$$= \frac{Area\ of\ monolayer\ trough}{Area\ of\ protein\ molecules}$$

$= 94.73 *10^{15}$ molecules

Therefore, maximum $94.73 *10^{15}$ molecules of ESAT-6 can interact with the membrane when the working concentration is 5μM.

The total surface area of a GUV with radius 15 μm is 2826 $μm^2$.

Thus, in 5μM ESAT-6 maximum number of molecules that can binds to GUV is $74.36*10^4$ molecules.

### Numerical simulation

We model the phagosomal membrane by simplifying it to a monolayered surface. The energy considerations for surface bending follow the Helfrich model[1], incorporating contributions from surface tension and active forces arising from ESAT-6 fibrillation. The overall energy functional is expressed as:

$$E = \int dA\,[2\kappa(C - C_0)^2] + \int dA\,\sigma + \int -d\boldsymbol{p}\,(F_R r + F_Z z)\,f(r,\theta,z(r)) \quad (1)$$

The three terms on the right-hand side of Eq. (1) designate the bending energy ($E_B$), surface tension energy ($E_\sigma$), and active forces energy ($E_F$), respectively. In the bending energy expression, $C$ represents the mean curvature of the surface, $C_0$ denotes the spontaneous curvature, $\kappa$ stands for the bending rigidity, and $dA$ represents the infinitesimal area element of the membrane. In the second term, σ signifies the surface tension. In the third term, $\boldsymbol{p} = (r,\theta,z(r))$, and $f\big(r,\theta,z(r)\big) = 1$ for the membrane region influenced by the fibrillation force; otherwise, $f\big(r,\theta,z(r)\big) = 0$ [2]. We note that, the membrane shape is described using the Monge representation where $\boldsymbol{p} = (r,\theta,z(r))$**.** The detailed description of the parameters used in the model are tabulated in Table 1.

We employ Monte-Carlo simulated annealing method to investigate the described model. This approach facilitates the identification of low-energy structures of the membrane under various parametric scenarios. In our simulation, we use surface triangulations method [3] to represents the discretized monolayered surface. Discrete version of the energy terms in Eq. (1) are used to calculate the total energy at each vertex of the membrane. Following the expressions given in [2], we calculate the mean curvature at vertex $i$ using $C_i = 0.5 \times (\mathbf{a}_i \cdot \boldsymbol{v}_i / \boldsymbol{a}_i \cdot \boldsymbol{v}_i)$ where $\boldsymbol{a}_i$ stands for the gradient of area and $\boldsymbol{v}_i$ for the gradient of volume computed with respect to the coordinates of the vertex $i$. The bending energy at vertex $i$ can now be recast as, $E_{B,i} = (C_i - C_0)^2 \times A_i/3$, where $A_i$ represents the total area of the triangles which share the same vertex. The spontaneous curvature ($C_0$) is induced at the vertices of the membrane where the ESAT-6 molecules bind. In the simulation, we consider a circular region around the central Z-axis with a specific radius (central region) on the flat membrane. The effect of spontaneous curvature on the initial flat membrane is depicted in Fig. supplementary. The energy due to surface tension is discretized as $E_\sigma = \sigma \left|\frac{\sum A_i}{3} - A_0\right|$, where $A_0$ is the reference area of the initial flat membrane. The surface tension energy accounts for the energy penalty due to change in surface area of the membrane.

In the model, we consider a resultant vertical force ($F_Z$) acting parallel to Z-axis and a resultant radial force ($F_R$) acting along the XY-plane towards the central Z-axis (Fig. 3i). This simplification proves meaningful when examining the complex network formed by the fibrils (Fig.3f-g), allowing us to anticipate the resultant vertical and radial forces. By adopting this specific force configuration, we aim to capture the essential aspects of the force components that impact the shape of the membrane. In the simulation, two force components are applied within the central region on the monolayered surface. This particular region has already undergone deformation as a result of spontaneous curvature. During force application, the significance of considering this specific region becomes evident. This is because the bound ESAT-6 molecules located in the central region of the surface undergo polymerization, leading to the formation of fibrils. These fibrils, in turn, exert forces on the membrane over this central region. The action of these two forces is the sequential formation of a tubular bud, caused by the vertical force component, and a spherical bud with a constricted neck, caused by the additional radial force component (see Main Text).

The Monte-Carlo simulation is performed by perturbing the surface and then pass it through Metropolis algorithm to find out the lowest energy configuration. The perturbation process involves two steps. At first a vertex is randomly displaced in a direction uniformly chosen from a cube $[-0.05l_0, 0.05l_0]^3$, where $l_0$ represents the average edge length of the triangles of the initial flat membrane. This step introduces random movements, contributing to the fluidity of the membrane. In second step, an edge on a rhombus undergoes flipping, which entails the removal of an edge shared by two triangles. Subsequently, the edge is reconnected in a manner that spans between the opposite vertices, which were previously unattached [2, 3]. This process serves the dual purpose of preserving mesh connectivity and introducing dynamic alterations to the membrane structure. We represent the initial flat membrane using $\sim 1000$ vertices, and then run $\sim 10^3$ Monte-Carlo steps in the simulation.

### Plausible fission mechanism of the spherical bud from the neck

Upon action of vertical and radial forces the membrane assumes a shape consisting a spherical bud attached to the flat membrane by a constricted neck (Fig.3j-q, Fz=10, Fr=4”). The fission of the spherical bud from the constricted neck, leaving behind the flat membrane, is a critical stage in membrane vesiculation. This process involves the separation of the spherical bud from the neck, resulting in the formation of two distinct entities. The constricted neck plays a pivotal role in mediating this fission event, acting as a bridge that undergoes a narrowing process before ultimately allowing the detachment of the spherical bud. Importantly, we believe that the phenomenon of line tension comes into play during this process. Line tension refers to the energy associated with the interface between the neck and the spherical bud, influencing the stability of the constricted region. The resulting phase separation of lipid molecules along the neck between the spherical bud and the remaining membrane is believed to be a consequence of line tension [4]. This separation is crucial for the redistribution and reorganization of lipid molecules to achieve a more energetically favorable configuration. The interplay of fission dynamics, line tension, and lipid phase separation contributes significantly to understanding the intricate processes governing vesiculation observed in phagosome.

Table 1: Description of the parameters used in the model

| Parameter | Description | Units |
| --- | --- | --- |
| $\kappa$ | Bending rigidity of the membrane | $80\ pN.nm$ [5, 6] |
| $C$ | Mean curvature of the membrane | $nm^{-1}$ |
| $C_0$ | Induced spontaneous curvature when ESAT-6 monomers bind to the membrane | $0.1\ nm^{-1}$ |
| $\sigma$ | Surface tension of the membrane | $0.5k_BT\ nm^{-2} - 1.0k_BT\ nm^{-2}$ [2] |
| $k_BT$ | Thermal energy | $4.1\ pN.nm$ [7] |
| $F_Z$ | Vertical force applied to the deformed membrane | $5\ pN - 10\ pN$ |
| $F_R$ | Radial force applied to the deformed membrane | $2\ pN - 4\ pN$ |

REFERENCES-
