## Supplementary data for "Real-time visualization reveals Mycobacterium tuberculosis ESAT-6 disrupts phagosome via fibril-mediated vesiculation"

### Extended Data

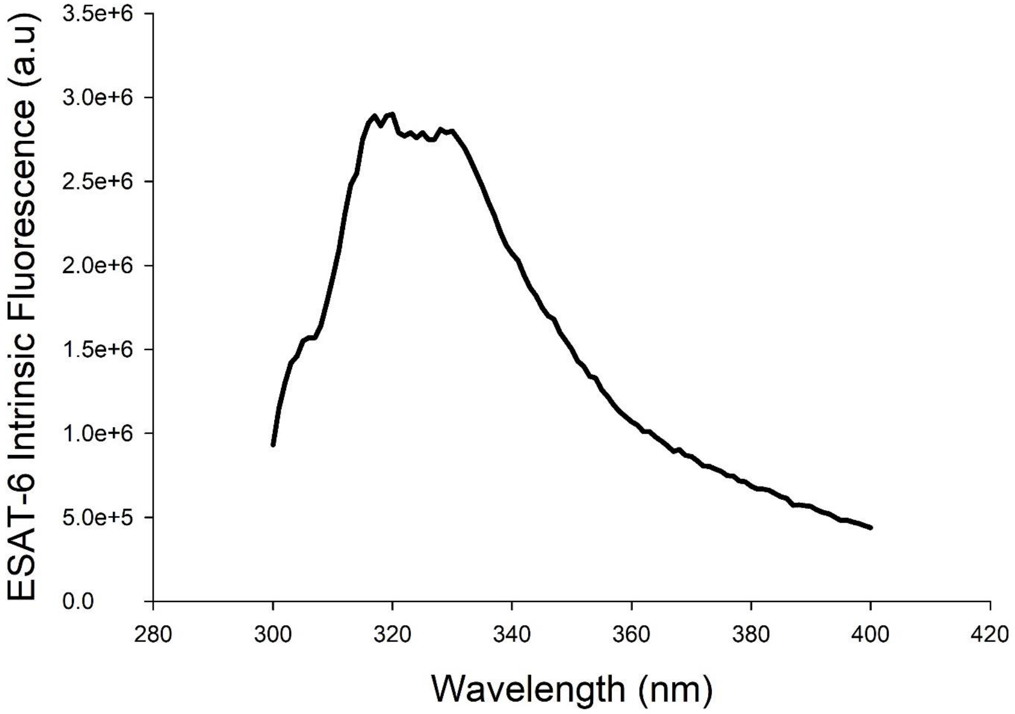


**Extended Data Fig.1 Emission spectra of intrinsic tryptophans of ESAT-6.** Emission spectra of the tryptophan residues of ESAT-6 show the maxima around 335nm. Experiments were carried out in PBS pH 5.5 at 25 °C. ESAT-6 concentration was kept constant at 5μM.

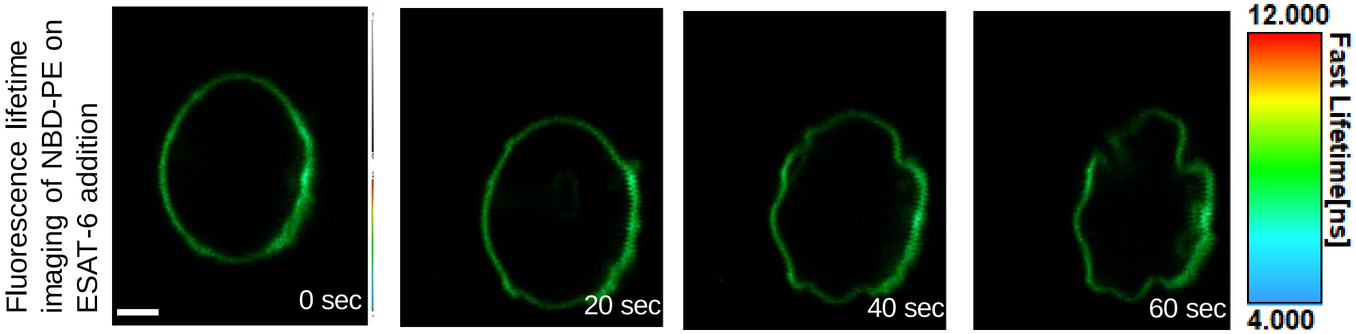


**Extended Data Fig.2.** Fluorescence lifetime imaging microscopy (FLIM) of Phagosomal membrane GUVs labeled with 0.1% NBD-PE (green) and immobilized using 0.03% PEG-Biotin incubated with 5μM unlabelled ESAT-6. Scale bar, 5μm.

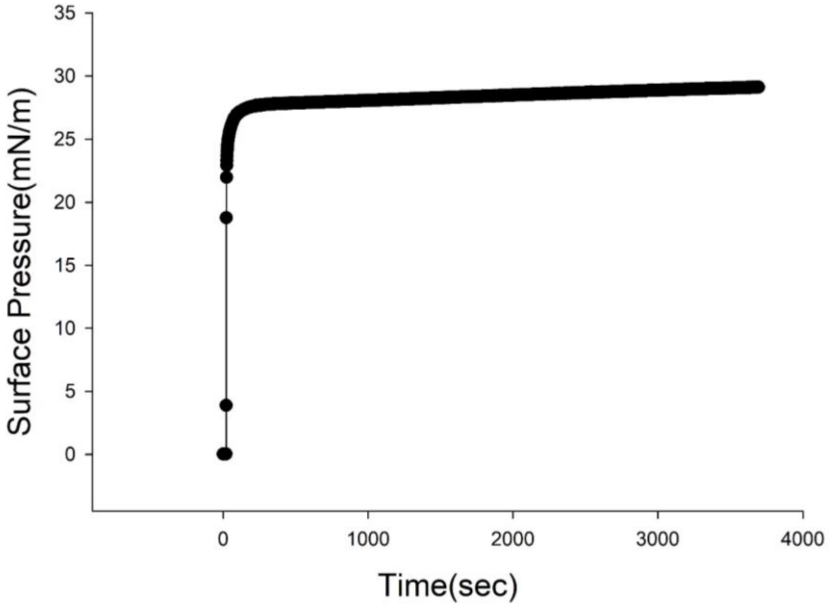


**Extended Data Fig.3. Surface Activity of ESAT-6**
Absorption kinetics of 5μM ESAT-6 as a function of time at the air/buffer interface. Each point corresponds to the mean of three different experiments. The subphase for all the experiments was performed in PBS(pH 5.5) and at 25C.

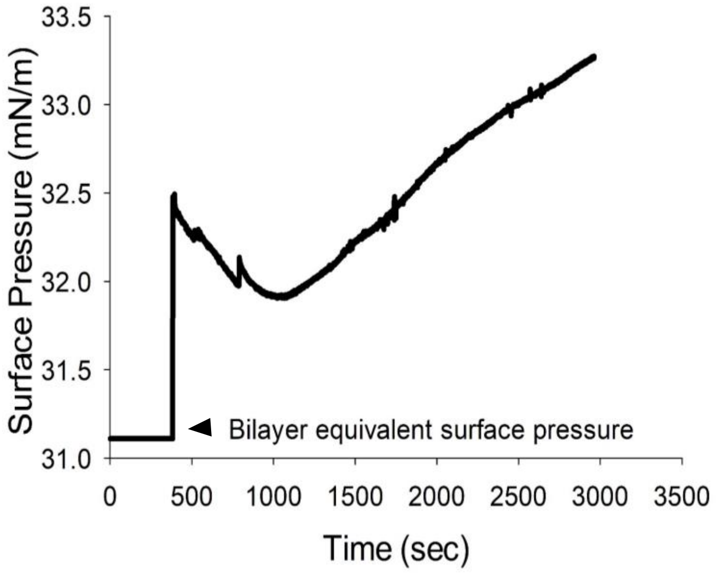


**Extended Data Fig.4. Interaction of ESAT-6 with the phagosomal membrane monolayer.**

Absorption kinetics (π-t) curves of ESAT-6 on the phagosomal membrane composition as a function of time at the air/buffer interface. ESAT-6 (5μM) was injected into the PBS(pH 5.5) beneath the monolayer compressed at an initial surface pressure π of 31.2mN/m. Each point corresponds to the mean of three different experiments.

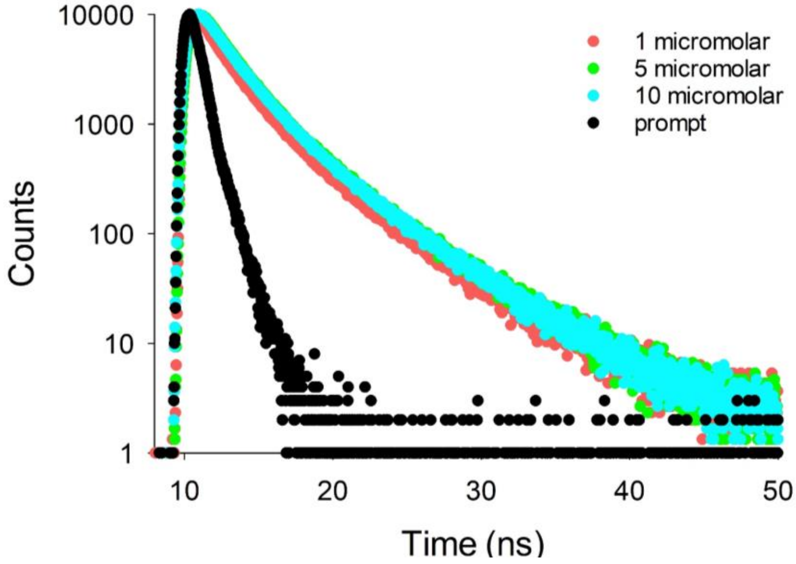


**Extended Data Fig. 5. Fluorescence decay plot of ESAT-6 after 30 minutes of incubation with the phagosomal membrane.**

Time-resolved fluorescence intensity decay plots of 1(red),5(green), and 10 μM(blue) of ESAT-6 respectively in the presence of phagosomal membrane mimic LUVs and measured immediately (30 minutes). The fluorescence decay plot of the instrument response function (IRF) is shown in black for both plots. All the measurement was carried out in PBS (pH 5.5) at 25°C. These are the representative curves from three independent experiments.

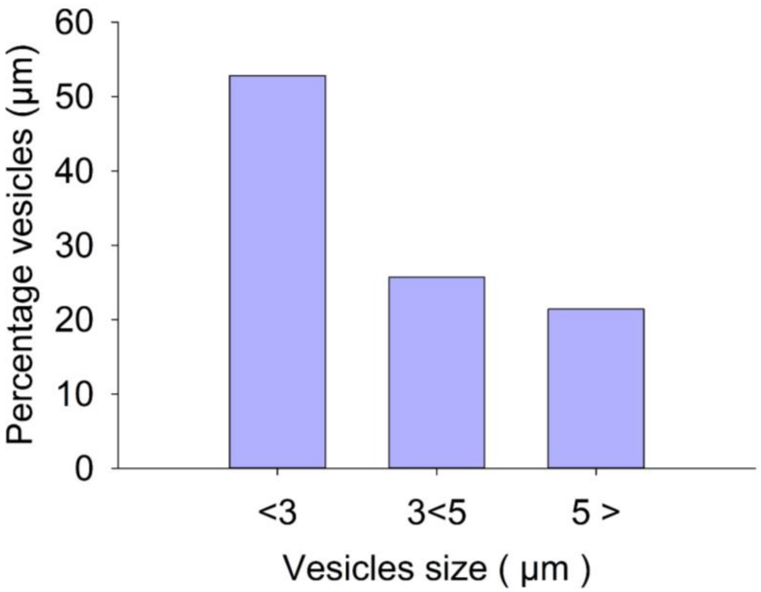


**Extended Data Fig.6.** Size distribution of the budded vesicles sizes (in µm) after the binding of 20µM ESAT-6.

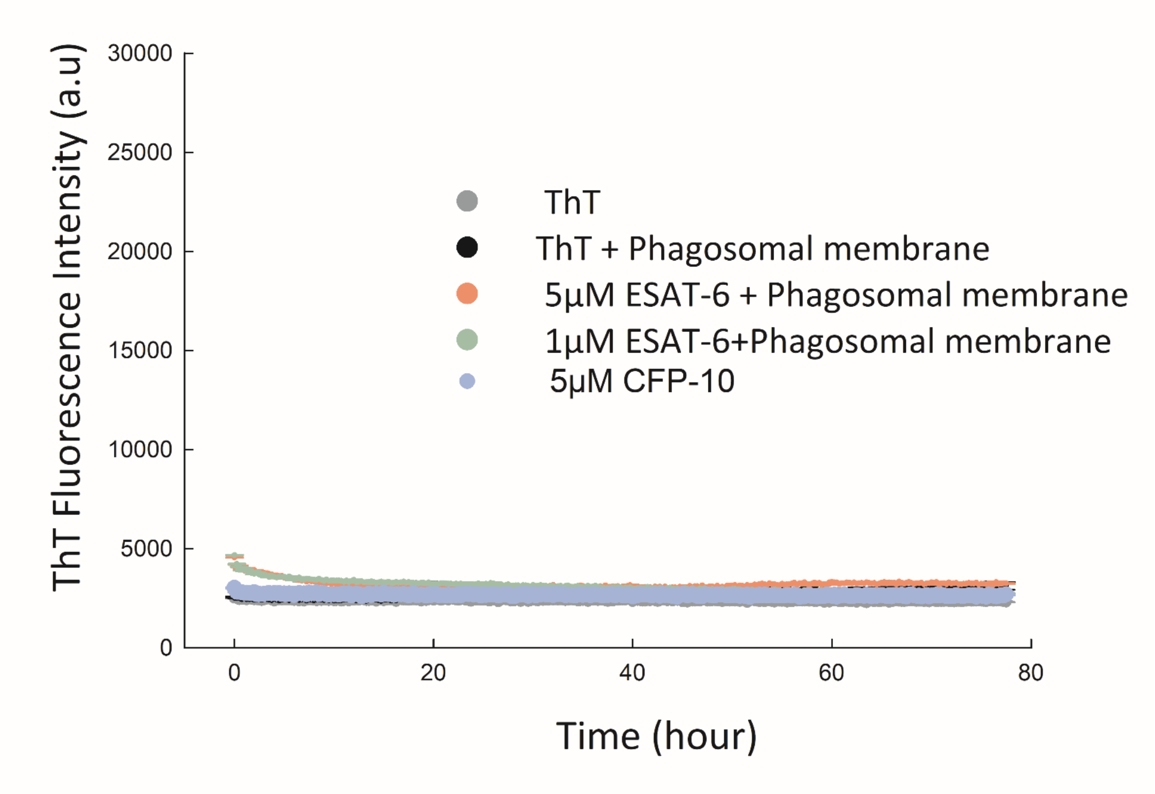


**Extended Data Fig.7. ThT fluorescence intensity of 1 & 5 µM ESAT-6 and CFP-10.**

ThT fluorescence assay of 1&5 µM ESAT-6 and 5µM CFP-10 as a function of time. Data points are shown as the means ± S.D. of three independent measurements. All the experiments are carried out on PBS pH 5.5 at 37°C**.**

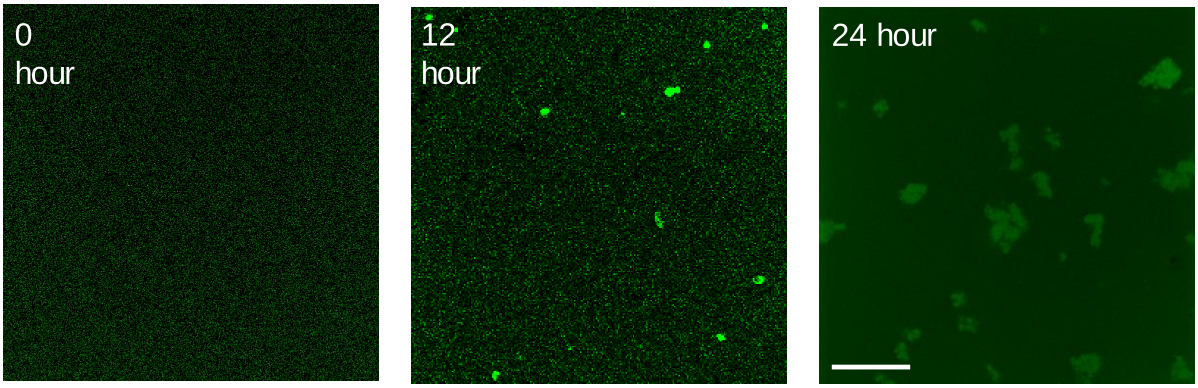


**Extended Data Fig.8. Aggregation of ESAT-6**
Time-lapse confocal imaging of 20μM FITC-ESAT-6 aggregation in PBS (pH 5.5) incubated and captured at ambient temperature over 24 hours. Scale bar, 25μm.

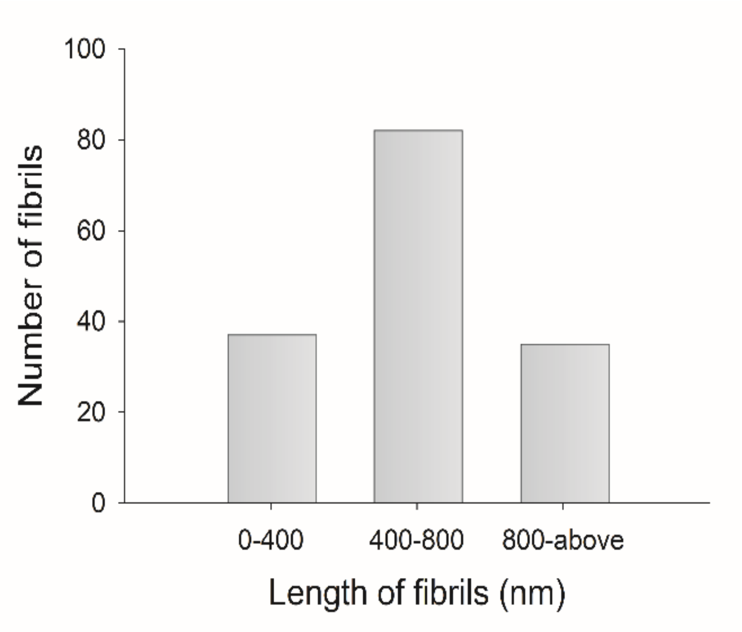


**Extended Data Fig.9. Length of ESAT-6 fibrils**
Number of ESAT-6 fibrils observed under the electron microscope of various lengths. (n=140nm fibrils measured through image-J)

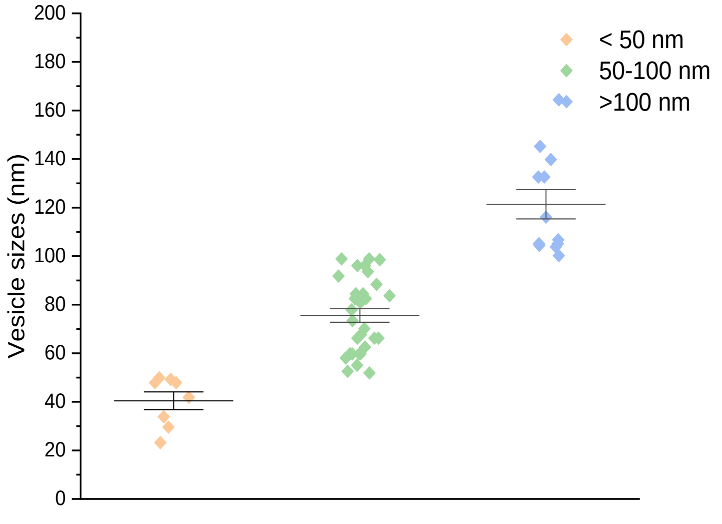


**Extended Data Fig.10. Size distribution of the vesicles after fission.** Statistics of vesicle sizes (in nm) after the vesiculation by ESAT-6.

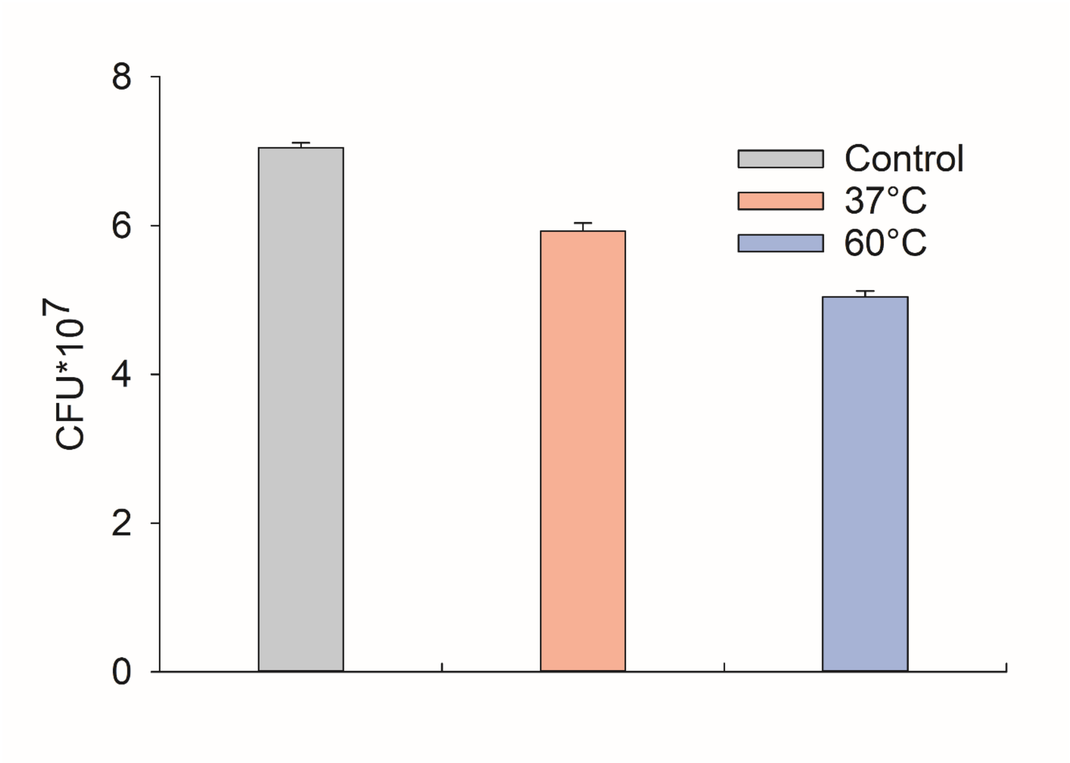


**Extended Data Fig.11. Quantification of mycobacterial viability.** Encapsulated *Mycobacterium smegmatis* viability was quantified by colony-forming units at different temperatures to mimic similar conditions as in electroformation. After treating the bacteria for 3 hours at those indicated temperatures, cells were lysed and log dilutions were spread on 7H11 agar plates. CFUs were counted on 4th day. Data quantitation wherever given is shown as Mean ± SD.

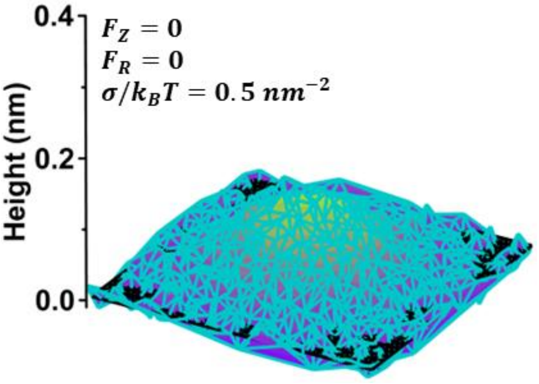


**Extended Data 12. Early shape transitions in membrane.** Membrane undergoes deformation induced by intrinsic curvature where $C_0 = 0.1\ nm^{-1}$ in the absence of the polymerization forces ($F_Z$ and $F_R$), providing insight into the intrinsic reshaping mechanism.
