## Supplementary material for "Real-time visualization reveals Mycobacterium tuberculosis ESAT-6 disrupts phagosome via fibril-mediated vesiculation": Movies: Supplementary Movie legends.pdf

**S1. Movie. ESAT-6 mediated membrane tubulation at low concentration.** Time-lapse confocal imaging of Phagosomal membrane GUVs labeled with 0.1% Rh-PE (red) and immobilized using 0.03% PEG-Biotin incubated with 5µM FITC-ESAT-6 (green). Scale bar, 5µm. Video accelerated by 2X.

**S2. Movie.** Time-lapse confocal imaging of Phagosomal membrane GUVs labeled with 0.1% Rh-PE (red) and immobilized using 0.03% PEG-Biotin incubated with 5µM unlabelled CFP-10 as a control. Scale bar, 10µm. Video accelerated by 2X.

**S3. Movie.** Time-lapse confocal imaging of Phagosomal membrane GUVs labeled with 0.1% Rh-PE (red) and immobilized using 0.03% PEG-Biotin incubated with 5µM unlabelled BSA as a control. Scale bar, 10µm.

**S4. Movie. Membrane budding at high concentrations(20µM) of ESAT-6.** Time-lapse confocal imaging of Phagosomal membrane GUVs labeled with 0.1% Rh-PE (red) and immobilized using 0.03% PEG-Biotin incubated with 20µM unlabelled ESAT-6. Scale bar, 25µm.

**S5. Movie. Real-time visualization of aspirated phagosomal membrane mimic.** Time-lapse fluorescence imaging of Phagosomal membrane GUVs labeled with 0.1% Rh-PE (red) aspirated by a negative pressure of 50pa and monitored over time.

**S6. Movie. Real-time membrane fission of aspirated tongue induced by ESAT-6 mediated.** Time-lapse fluorescence imaging of Phagosomal membrane GUVs labeled with 0.1% Rh-PE (red) aspirated by 50pa force and incubated with 20µM ESAT-6.
